# Cell type transcriptomics reveal shared genetic mechanisms in Alzheimer’s and Parkinson’s disease

**DOI:** 10.1101/2025.02.17.638647

**Authors:** Anwesha Bhattacharya, Edward A. Fon, Alain Dagher, Yasser Iturria-Medina, Jo Anne Stratton, Chloe Savignac, Jack Stanley, Liam Hodgson, Badr Ait Hammou, David A Bennett, Danilo Bzdok

**Affiliations:** Department of Biological and Biomedical Engineering, McGill University; Montréal, Canada; Mila - Quebec Artificial Intelligence Institute; Montréal, Canada; Department of Neurology and Neurosurgery, Montreal Neurological Institute (MNI), McGill University; Montréal, Canada; Department of Psychology, MNI, McGill University; Montreal, Canada; McConnell Brain Imaging Centre (BIC), MNI; Montreal, Canada; Ludmer Centre for Neuroinformatics and Mental Health; Montreal, Canada; Quantitative Life Sciences, McGill University; Montreal, Canada; School of Computer Science, McGill University; Montreal, Canada; Rush Alzheimer’s Disease Center, Rush University Medical Center; Chicago, USA; The Neuro, MNI, BIC, McGill University; Montreal, Canada

## Abstract

Historically, Alzheimer’s disease (AD) and Parkinson’s disease (PD) have been investigated as two distinct disorders of the brain. However, a few similarities in neuropathology and clinical symptoms have been documented over the years. Traditional single gene-centric genetic studies, including GWAS and differential gene expression analyses, have struggled to unravel the molecular links between AD and PD. To address this, we tailor a pattern-learning framework to analyze synchronous gene co-expression at sub-cell-type resolution. Utilizing recently published single-nucleus AD (70,634 nuclei) and PD (340,902 nuclei) datasets from postmortem human brains, we systematically extract and juxtapose disease-critical gene modules. Our findings reveal extensive molecular similarities between AD and PD gene cliques. In neurons, disrupted cytoskeletal dynamics and mitochondrial stress highlight convergence in key processes; glial modules share roles in T-cell activation, myelin synthesis, and synapse pruning. This multi-module sub-cell-type approach offers insights into the molecular basis of shared neuropathology in AD and PD.

## Introduction

Alzheimer’s disease (AD) and Parkinson’s disease (PD) are two of the most prevalent disorders in today’s aging societies ^1,2^. There has been intensive research with the grand aim of altering and ultimately halting the course of these diseases. Despite educated forecasts predicting significant advances by this decade ^3^, AD and PD remain challenging to unravel. This difficulty is compounded by a historically entrenched dichotomy that has limited transfer of research insights from one disease to the other. AD and PD are considered distinct entities due to differences in primary brain regions affected, age of onset, clinical progression, and treatment response. PD is notably responsive to therapeutics that do not affect cognition ^4^, and AD is without any “hard-currency” therapeutic to date ^5^.

This dichotomy has been reinforced by genomics and polygenic risk studies which show minimal to no overlap of genes between AD and PD ^6,7^. Indeed, aggregating prior findings, a *Neuron* review recently concluded, “There is intriguingly little overlap between the risk genes for AD and PD, providing genetic evidence for different disease onset and progression mechanisms” ^8^. However, over half of PD patients show aggregates of tau ^9^ and around 30% of PD patients develop cognitive impairment with many going on to dementia ^10^. Conversely, AD pathology and Lewy bodies co-occur more frequently than by chance, with Lewy bodies associated with fluctuating cognitive decline ^11–13^. Further, the substantia nigra in AD can harbor tangles which are symptomatic of parkinsonism ^14^. These hints raise the possibility of shared disease mechanisms between AD and PD at the molecular level ^15,16^.

Most studies investigating the genetic basis of the neuropathological overlap between AD and PD have predominantly employed approaches that center on univariate approaches focussing on single genes. Prominent among these are genome-wide association studies (GWAS), and differential gene expression (DGE) analyses ^17–19^. However, gene expression occurs within tightly regulated environments where gene products interact in highly combinatorial ways ^20–23^. Thus, the pathogenesis of neurodegenerative diseases is likely driven by molecular dysregulation within gene networks rather than isolated gene anomalies ^24,25^. Further, the effects of dysregulated genes are not confined to a single type of cell. For example, there is growing evidence, in both AD and PD, that the protective functions of glial cells can lead to a vicious cycle of reactive oxygen species (ROS) production that exacerbates neuronal death cascades, which hyper-excites additional glial cells ^26–28^. Thus, this pathological process becomes a candidate for being shared across both diseases.

Given this complexity, it is crucial to account for the interaction between genes and cell type heterogeneity when studying these diseases. Recent advances in single-nucleus RNA-sequencing (snRNA-seq) have provided the ability to resolve transcriptional changes with high cell-type specificity. For example, large-scale single-nucleus transcriptomic studies in AD ^29,30^ and PD ^31,32^ have reported significant disease-associated transcriptional changes that are highly specific to certain cell type populations.

In the present investigation, we systematically revisited the problem of identifying candidate molecular mechanisms that might overlap between AD and PD. Our unique approach focused on studying the concerted effects of genes at a sub-cell-type granularity. Comprehensive datasets from AD (70,634 nuclei from 48 postmortem brains) ^30^ and PD (340,902 nuclei from 15 postmortem brains) ^33^ allowed us to leverage advanced machine-learning techniques ^34^. Enabled by a supervised multivariate model ^35^, we extracted several biologically interpretable gene modules within cell types from the AD and PD transcriptomes (16,936 protein-coding transcripts). This method combined the latent structure discovery strength of techniques like tSNE, UMAP, and variational autoencoders ^36^ while simultaneously being aware of valuable contextual information — the disease state. Additionally, by focusing our inter-disease comparative analysis solely on gene module compositions, we circumvented potential batch effects of combining the raw transcriptomic datasets that could have compromised our analysis. By linking our gene programs to a catalog of biological pathways in the brain, we discovered shared cellular basis of neurodegeneration. We further validated our main findings and conclusions using an independent AD and PD dataset pair ^32,37^. Thus, our integrated bottom-up approach comparing multiple modes of genetic interactions sheds new light on the molecular mechanisms of AD and PD.

## Results

### Genetic correlation between AD and PD uncovered in multiple gene modules across major brain cell types

We explored the possibility of molecular overlap in AD and PD brains through transcriptional alterations captured in gene modules (groups of co-expressed genes). Our main analyses were conducted on a snRNA-seq AD dataset ^30,38^ (70,634 nuclei across 8 major cell types; referred to as ROSMAP-AD) and a PD dataset ^33^ (340,902 nuclei across 11 cell types; referred to as Kamath-PD). Our analytical framework employed partial least squares discriminant analysis (PLS-DA) to gain an overview of 16,936 protein-coding transcripts (Fig. 1; see Methods). In either AD or PD, we fitted cell type level PLS_cell_ models across nuclei from all donors in the dataset. This yielded gene programs (gene modules) as latent projections of gene expression matrices that assisted in distinguishing cells of patients from controls ^35^. Comparative assessments of these thus-derived gene modules highlighted shared genetic mechanisms between AD and PD that were more stable than expected by chance (Fig. 2A).

**Figure 1.**
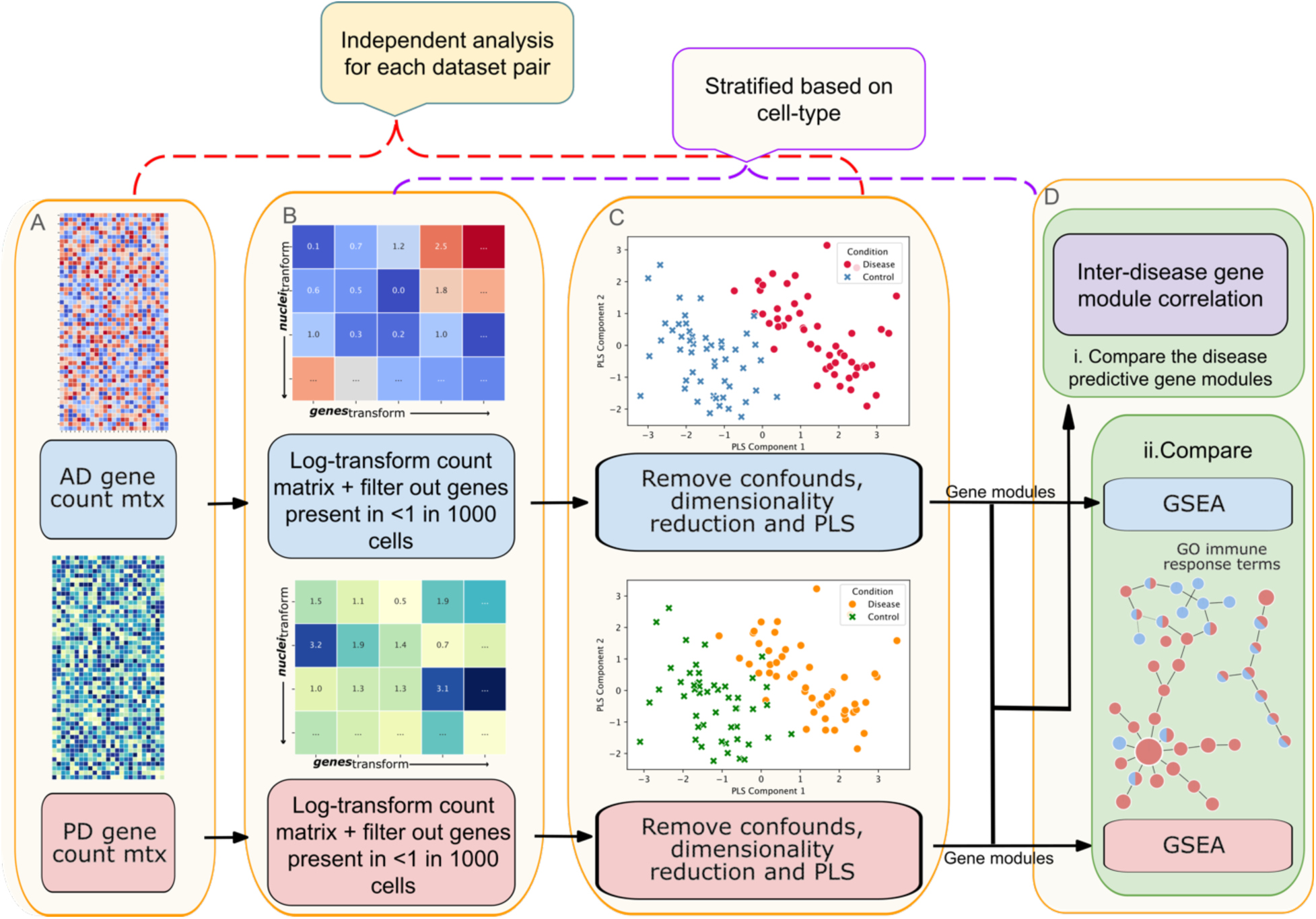
Workflow diagram to test for AD-PD overlap: bottom-up approach. Overview of workflow. **(A)** Single nucleus RNA sequencing (snRNA-seq) datasets for AD and PD were downloaded from public databases. Filtering and quality control of recordings were already done by source authors, as were cell type annotations. **(B)** Pre-processing step. Performed independently for each cell type, this step followed recommended guidelines for data transformation, removed lowly expressed genes, and corrected disease vs control class imbalance. **(C)** Gene module identification step. PLS discriminant analysis was performed per cell type to extract weighted gene lists, referred to as gene modules, that were disease predictive. **(D)** Comparative analysis. We aggregated the results from the two analysis arms and looked at the level of overlap using parallel methods – i. direct correlation of cross-disease gene module pairs and ii. gene set enrichment analysis to identify overlapping biological processes, molecular functions and cellular components. We discover significant molecular similarities between AD and PD across cell types. PLS, Partial least squares; AD, Alzheimer’s disease; PD, Parkinson’s disease; GSEA, gene set enrichment analysis; GO, gene ontology.

**Figure 2.**
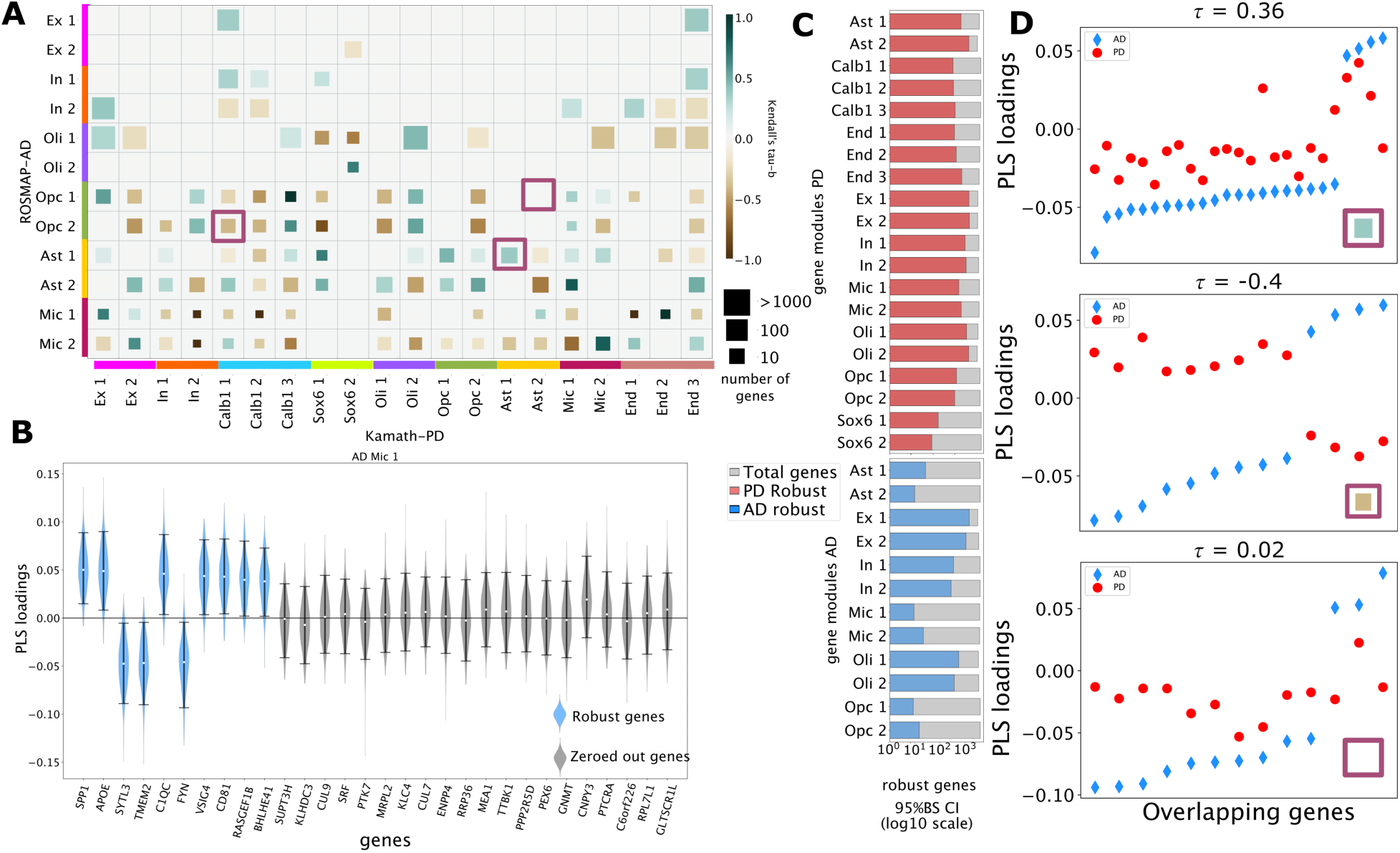
Convergence of disease mechanisms in Alzheimer’s and Parkinson’s disorders across cell-types when zoomed in on disease-predictive gene groups. We probed two snRNA-seq datasets (ROSMAP-AD, Kamath-PD) to explore the genetic overlap between AD and PD. By training 15 PLS models (one for each cell type in AD or PD), we extracted latent representations of gene expressions (gene modules) that maximized the separation between disease and healthy nuclei. **(A)** The coloured squares represents kendall’s tau-b (*τ*_b_), quantifying the degree of association between AD-PD gene modules. Each pairwise *τ*_b_ is statistically significant exceeding 2.5/97.5% CI based on label shuffled permutation test (n=1000). Darker colors represent stronger genetic associations, indicating similar ranking trends of robust genes in the module pair. Square size represents number of robust genes shared between the two modules. Robust genes were assessed based on a 500-iteration bootstrap resampling scheme. Each colored bar (left vertical and bottom horizontal) represents a unique PLS model fitted on a cell type pre-identified in the dataset. The wide range of correlation strengths suggested that AD and PD shared significant genetic similarity spanning the subcellular landscape. **(B)** An illustration of the robustness assessment of genes in a gene module. The top 30 genes with the highest empirical predictive weights from the first PLS component in AD microglia are shown. Violins represent the loading distribution of genes derived from a bootstrap resampling scheme (n=500). Robust genes (blue) have non-zero predictive weights (2.5/97.5% confidence interval). Genes that do not meet this criterion are greyed out. APOE, a gene strongly associated with AD, was highlighted as a major disease predictor in this component. **(C)** The bars show the number of genes with robust disease predictive weight for each gene module. The colored portion of the bars denotes the number of robust genes in a module, while the grey region shows the total analyzed genes for that cell type. Different modules have different numbers of robust genes. **(D)** Select gene module correlations are visualized. Top, an example of a strong positive *τ*. Center, strong negative *τ*. Bottom, weak *τ*. Non-zero overlapping genes between a chosen gene module pair are represented on the x-axis. Ast, Astrocyte; Ex, Excitatory neuron; In, Inhibitory neuron; Mic, Microglia; Oli, Oligodendrocyte; Opc, Oligodendrocyte precursor cell; PLS, Partial least squares.

As a first step, we explored the transcriptomes of AD and PD individually to identify disease-specific modeling parameters. For each cell type, the number of modules that best distinguished diseased from healthy cells in unseen brain samples was treated as an hyperparameter in our model selection. We used a 10-fold cross-validation scheme on the stratified dataset containing nuclei from one cell type to estimate this. In AD, 2 gene modules per cell type were determined as optimal for all 6 cell categories (2 categories, ependymal cells, and pericytes, were removed from further analysis due to low nuclei count). In PD, the optimal number of gene modules was also determined to be 2 for each of the 9 major cell types (2 categories, macrophages, and ependymal cells, were removed from further analysis due to low nuclei count), except for endothelial cells (3 modules) and CALB1 dopaminergic neurons (3 modules). Each gene module, on its own, represented combinations of several genes whose co-expression signature was associated with a disease state (AD vs. control or PD vs. control). The statistical significance of a module was assessed by comparing its empirical disease prediction performance to a null distribution of performance metrics derived from a label-shuffling permutation procedure (one-sided p-value < 0.05). As an illustration, Fig. 2B visualized the PLS_Mic_ loadings for the first gene module from ROSMAP-AD microglial cells.

All thus chosen PLS_cell_ models (6 AD and 9 PD models), exhibited robust above-chance out-of-sample accuracy in differentiating disease samples from control, as evidenced by the area under the receiver operating characteristic curve (AUROC) scores (fig. S1A). Unbiased classification accuracy was estimated based on a patient-partitioned cross-validation scheme (see Methods). In the AD vs control group contrast, the predictive power measured by AUROC ranged highest for microglia (AUROC: 0.66 ± 0.06 std across partitions) to lowest for oligodendrocyte precursor cells (OPCs) (0.56 ± 0.08 std across partitions). For the PD vs control group contrast, the highest AUROC was for endothelial cells (0.89 ± 0.13 std across partitions), and the lowest was for excitatory neurons (0.69 ± 0.35 std across partitions). This strongly supported the role of gene modules in directly informing the disease phenotype across all examined cell types and conditions.

We further inspected the 16,936 gene coefficients in each PLS_cell_ module (12 AD modules, 20 PD modules) that captured the contribution of a gene within the module. Specifically, using a bootstrap (BS) resampling technique (see Methods), we assessed which gene effects were statistically robust (zero not included in the 2.5/97.5% confidence interval (CI) of the BS distribution of each gene), and thus, robustly affected prediction outcomes. Each gene module yielded a variable set of genes distributed across the transcriptome (Fig. 2C). We noted that most gene modules in both AD and PD retained tens to hundreds of robust genes from the 16,936 genes examined. In AD, astrocytes, microglia, and OPC modules had the smallest number of robust genes (10 to 100 per module). In the PD dataset, SOX6 gene modules had the most specific subset of robust genes (∼100 per module).

To explore the characteristics of the derived modules, we investigated if they represented cellular subpopulations which would be highlighted in a typical clustering algorithm. For a cell type, we assigned each observed nucleus to exactly one of its modules based on the component harboring the maximum PLS_cell_ score. In doing so, we were able to visualize the distribution of the gene modules assigned to the nuclei in a two-dimensional embedding space. The embedding space was derived using PHATE ^39^ from the ambient gene space containing all nuclei from the cell type (fig. S2). No clear clustering among the components was observed. Thus, we concluded that our gene modules did not necessarily correspond to cellular subtypes, but likely different functional programs or biological states within a given cell type; that is, any given cell belonging to a type can exhibit several distinct expression programs to various continuous degrees.

Importantly, the samples from AD and PD datasets were not merged. Instead, all analyses, so far, was conducted independently in AD and PD. In so doing, we bypassed certain conceivable batch effects faced by many alternative methods that aim to directly combine different RNA-seq transcriptomes into a single composite dataset ^40^. We also noted that our data-driven approach examined the entire transcriptome without prior assumptions about specific genes generally associated with AD or PD. Moreover, our approach could assign a single gene as relevant to multiple modules. In contrast, previous work using gene clustering or co-expression networks (which also aim to identify gene groups) often overlooked interactions between gene products across different biological pathways ^25^. Additionally, they frequently started by seeding the networks with apriori-determined genes to build interaction graphs ^41,42^. Predefined genes, typically chosen based on a single-gene disease perspective, have limitations outlined earlier.

We subsequently moved to our comparative analysis between AD and PD. To quantify the coupling between pairwise gene modules, we used Kendall’s tau-b (*τ*_b_) metric. It calculated the degree of similarity between two vectors of PLS_cell_ predictive weights (not the gene expression measurements; Fig. 2D) of the overlapping robust genes from an AD-PD gene module pair. By comparing the empirical correlation strength with a permutation-derived null distribution (obtained by correlating modules derived from a label-shuffled dataset), we identified the module pairs with significant associations (empirical *τ*_b_ more extreme than 2.5/97.5% CI of the null distribution). To further test the generalizability of our comparison model, we divided our original datasets from AD and PD into two random non-overlapping subset pairs (split-half test; see Methods) and observed strong correlations (0.92±0.02 std.) across different realizations of the under-sampled but otherwise identical analysis (fig. S1B). This indicated that the cross-disease gene module correlations are reproducibly detected even with fewer transcript samples.

As the most important results of our investigation so far, we noted strong and reliable associations among inhibitory neuron modules, among excitatory neuron modules, and among oligodendrocyte modules from AD and PD (Fig. 2A; table S1). Strikingly, every gene module from one disease featured some degree of overlap with at least one gene module from the other disease, within or between cell types. Across all pairwise combinations, the strongest correlation was found between the first oligodendrocyte module (represented as Oli 1) module from AD and the second oligodendrocyte module (Oli 2) from PD (represented as Oli 1_Oli 2; *τ*_b, abs_= 0.44, number of shared genes = 1351). We noted that in a parallel analysis involving DGE (described in a later section), a cross-disease cell-level comparison (based on the log-fold change between disease and control) reported a maximum association of *τ*_b, abs_= 0.20. In contrast, our native whole-transcriptome analysis described multiple sets of module combinations as having high degrees of similarity.

In addition to Oli 1_Oli 2, several other pairs (Oli 1_Ex 1, Oli 1_Ex 2, In 2_Ex 1) featured *τ*_b, abs_ > 0.30, and shared over 500 robust genes. Notably, dopaminergic neurons from midbrain tissue samples in PD, CALB1, and SOX6, also showed significant similarities with AD-critical gene programs from neurons, oligodendrocytes, and OPCs (In 1_Calb1 1, *τ*_b, abs_ = 0.32, number of shared genes = 100; Opc 2_Calb1 1, *τ*_b, abs_ = 0.41, number of shared genes = 13). On the flipside, inhibitory neuron module pairs between AD and PD showed the least overlap between each other. This suggests potentially different genetic contributions of these cell types to AD and PD disease mechanisms. Congruently, AD-derived astrocyte, microglia, and OPC gene modules demonstrated strong correlations with most PD modules. Particularly, Mic 1 from AD featured correlation strengths of *τ*_b, abs_ > 0.67 with both Mic 1 and Mic 2 from PD (p-value < 0.05, number of shared genes > 10). Similarly, Ast 2_Ast 2 featured *τ*_b, abs_ = 0.62 with 23 shared genes. Thus, these findings located significant sub-cell level genetic associations between AD and PD-relevant gene modules with the strongest associations among combinations of neuron and oligodendrocyte modules from AD and PD.

To replicate our primary findings, we examined the transcriptomic space using a different pair of snRNA-seq datasets related to AD (Seattle-AD) and PD (Smajić-PD), with independent cohorts. We repeated all main analyses and derived sub-cell-level gene modules from these datasets (detailed cohort and sample description in Methods). As before, our external validation analysis also revealed significant gene module overlaps between AD and PD, scattered across different cell types (fig. S3A; table S2). Oligodendrocyte module pairs from AD and PD, once again, took center stage with strong associations between each other as well as with modules from neurons, astrocytes, and microglia. Strong significant associations were also observed between different combinations of neuron and glial cell-derived modules. As in the primary analysis, inhibitory neuron-derived modules showed sparse similarities between AD and PD. Overall, the external validation of shared genetic signatures between AD and PD provided evidence that the primary observations were not merely a technical artifact of the specific primary transcriptomic samples, the participant cohorts, or the brain regions chosen.

Taken together, these findings revealed molecular overlap between AD and PD at sub-cell resolution. The gene module associations extended between and across distinct AD-PD cell types. The degree and specificity of these overlaps varied among cell types, with oligodendrocyte and neuron-based gene module combinations in AD and PD signaling the strongest similarities. Our external validation experiments replicated these core findings in independent datasets, further substantiating our conclusions regarding the shared genetic architecture between these neurodegenerative diseases.

### Cell type-specific gene modules reveal GWAS-linked genes as key predictors of disease

We contextualized our gene modules post-hoc to understand their relationship with known gene variants from genomic studies. Drawing from the most recent GWAS that reported AD ^43^ or PD ^44^, we looked at 164 genes (table S3; see Methods) and located them within our gene modules. Most GWAS genes showed significant disease-predictive loadings in at least one gene module (Fig. 3). The top gene module harboring the most GWAS genes in AD was Ex 1 with 25.6% of AD GWAS genes present, followed by Ast 1 with 24.3% GWAS genes present. In PD, the top modules were Oli 1 with 58.9% PD GWAS genes followed by Ex 2 with 42%. Moreover, we found clear cell type localization of these genes within this modular framework. For example, APOE, a broadly accepted AD risk gene, showed robust predictive loadings in the Ast 1 and Mic 1 derived from AD. Similarly, LRRK2, one of the major PD risk genes, was implicated in distinct PD-related gene modules. The strongest effect was observed in PD Mic 2. This cellular localization further corroborated our gene module modeling approach.

**Figure 3.**
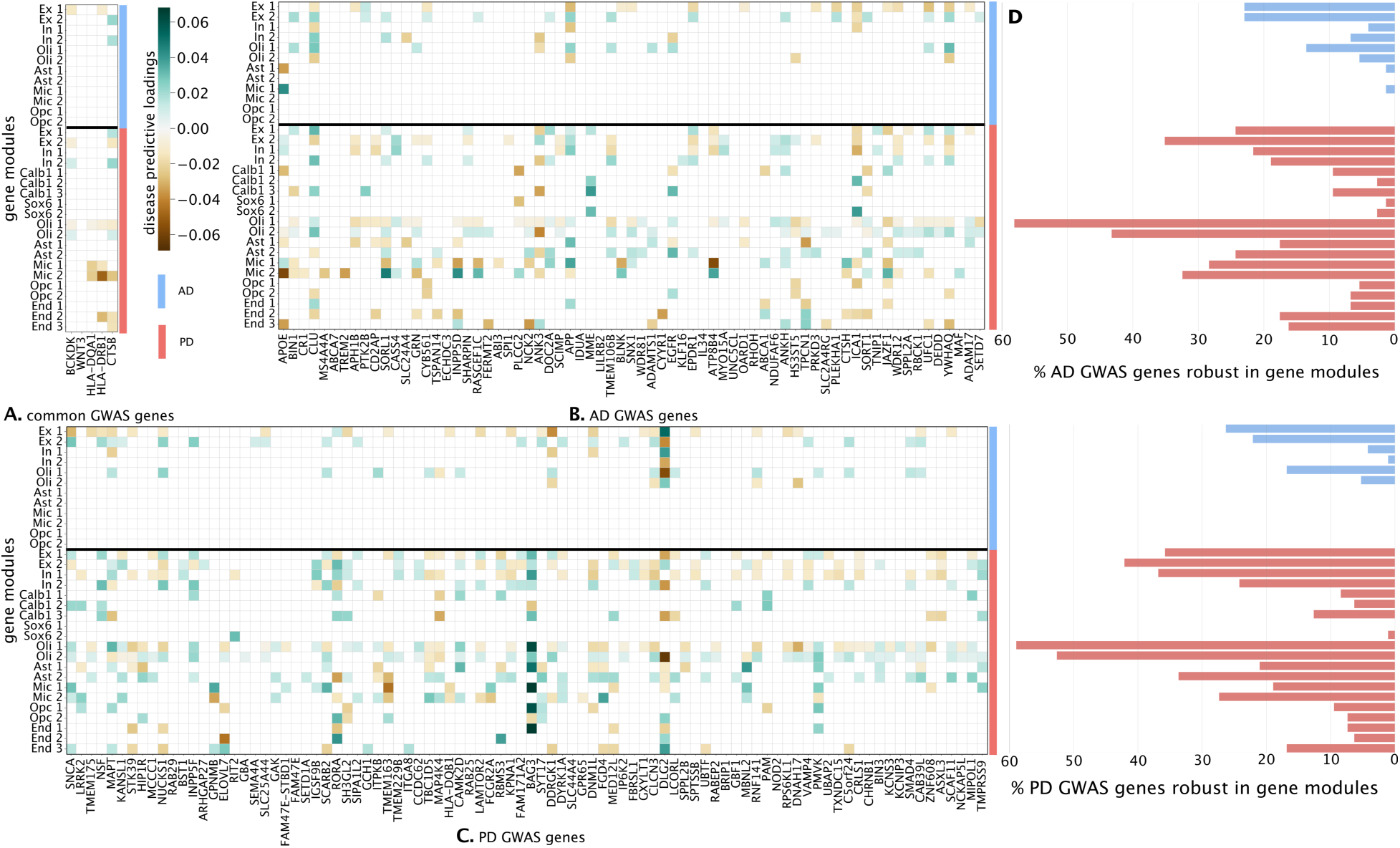
Cell type-specific gene modules reveal GWAS-linked genes as key disease predictors. We mapped the contribution of 164 risk genes (combined from the largest AD GWAS study and PD GWAS study) within our gene modules. The color saturation represents the strength of the predictive weight of a gene in a module as deduced by the PLS model (robust loading according to bootstrap iteration scheme passing 2.5/97.5% CI). **(A-C)** Genes grouped based on the nominating disease. Group (A) harbors 5 common genes nominated independently in both AD and PD. (B) Shows the genes implicated in AD GWAS. (C) Shows genes implicated in PD GWAS. **(D)** The bars on the right summarize the connection of a module with previously implicated risk genes. The lengths indicate the percentage of risk genes with robust loading in the module (blue = AD modules, red = PD modules). Cell-level localization of several key risk genes was observed. For example, APOE, a major AD risk gene, showed strong predictive loading in AD microglial and astrocyte modules. Similarly, SNCA had strong predictive signals in PD-based oligodendrocytes, microglia, excitatory neurons, and CALB1 dopaminergic neurons. This corroborated the gene compositions in our modules with previous complementary research. Notably, as a novel find, several genes which were implicated as being AD relevant in GWAS, had robust predictive loadings in PD gene modules and vice versa. For example, APOE, APP, and other AD risk genes had robust predictive weights in PD modules. Likewise, SNCA, MAPT, and other PD risk genes had robust predictive weights in AD modules. Ast, Astrocyte; End, Endothelial; Per, Pericyte; Ex, Excitatory neuron; In, Inhibitory neuron; Mic. Microglia; Oli, Oligodendrocyte; Opc, Oligodendrocyte precursor cell; GWAS, Genome-wide association study.

Next, we analyzed the effects of GWAS genes from one disease within gene modules linked to the other neurodegenerative disease. In other words, we investigated whether AD risk genes were highlighted in any PD modules and vice versa. Among the AD GWAS genes, top genes, including APOE, APP, BIN1, and CLU (based on p-value from GWAS ^43^) had robust disease predictive loadings in gene modules associated with PD. Concretely, APOE had the strongest predictive weight in PD Mic 2 (gene loading = -0.06, max absolute loading for any gene in this module was 0.07, absolute rank of this gene = 23), APP’s maximum loading was in PD Mic 1 (loading = 0.04, max_abs_ = 0.07, rank = 262), BIN1 was found in PD Calb1 2 (loading = -0.02, max_abs_ = 0.04, rank = 1626), and CLU had the strongest robust weight in PD Oli 2 (loading = 0.03, max_abs_ = 0.08, rank = 405). It is important to note that these genes were not the highest-ranked within their modules. In other words, they were not the primary disease-indicative genes (rank = 1). Instead, these genes likely played supporting roles that become apparent only within the context of gene modules.

Conversely, we made similar observations for trusted PD GWAS genes ^45^ within our AD modules. SNCA had strong predictive loading in AD Ex 1 (-0.03, max_abs_ = 0.07, rank = 397), MAPT in AD In 1 (-0.02, max_abs_ = 0.07, rank = 979), and TMEM175 in AD Ex 1 (-0.02, max_abs_ = 0.07, rank = 1869). Collectively, we observed that the bona fide GWAS genes not only tracked the disease they were implicated in, but known GWAS hits also proved relevant in gene modules associated with the other disease.

Notably, while the genes implicated in AD had very clear cell type localizations, the risk genes associated with PD tended to be distributed across modules implicating multiple cell types, with particularly high effect sizes in neuronal modules. This observation aligned with previous studies showing that PD risk loci are not confined to specific cell types. Instead, they are associated with broad cellular processes observable across multiple cell types ^46^.

In our external validation analysis, GWAS genes had robust disease-predictive loadings across the gene modules derived from Seattle-AD and Smajić-PD. Indeed, all observations highlighted in our primary dataset could be replicated (fig. S3 B-E). First, the genes tracked disease-corresponding modules; Microglia and astrocyte AD modules faithfully tracked APOE (interestingly, we observed robust predictive loading for APOE in OPC modules ^30^). Similarly, the PD GWAS genes, such as SNCA and LRRK2, were tracked by PD predictive modules. Second, as with the initial dataset pair, we observed strong representations of genes within gene modules associated with the disease opposite to the one implicated by GWAS, showcasing again the cross-disease genetic overlap.

We have thus successfully situated pre-established GWAS genes within the scope of gene modules. We have also identified significant effects of these genes beyond the specific disease in which they were originally reported in scientific literature. For example, not only was the impact of APOE most significant in AD microglia and astrocyte-specific modules, but we also observed robust disease-predictive signatures for APOE in PD modules corresponding to microglia, astrocytes, and CALB1 DA neurons. These results were consistently replicated in a validation test thus confirming our primary findings.

### Overlapping biological functions between AD and PD gene modules

To uncover the biological relevance of the gene modules, we conducted a comprehensive gene set enrichment analysis (GSEA) across the modules identified for each cell type in our latent factor analysis. We carried out this contextualization separately for AD and PD, analogous to our previous analysis steps. Notably, in our analysis, a single gene can contribute significantly to a disease via multiple modules within the same cell type; this, in turn, enabled us to capture its effect on complementary, co-regulated pathways. We screened the widely relied upon gene ontology databases (GO 2023) corresponding to three complementary domains - biological processes (BP), molecular functions (MF), and cellular components (CC). We focused on GO, as collectively, they cover the largest fraction of the genome. Each gene module was enriched in several significant GO terms (Benjamini-Hochberg corrected FDR<0.05). Across all gene modules, GO enrichment for AD highlighted 196 BP, 35 MF, and 73 CC terms. For PD, we obtained 561 BP, 103 MF, and 163 CC terms.

We next examined if any enriched terms overlapped between AD and PD. In total, 122 BP (out of 27,993 terms), 26 MF (out of 11,271 terms)., and 65 CC (out of 4,039 terms) terms were shared between AD and PD (Fig. 4A). The highest number of GO term overlaps emerged in neuron-neuron AD-PD module pairs and pairs involving oligodendrocytes and oligodendrocyte precursor cells (>160 terms per module pair, BP, MF, and CC combined; see Fig. 4B, also note Fig. 4C horizontal axis). Additionally, both AD and PD microglial gene modules shared on average 20 common term hits with other gene modules. In contrast, module pairs involving astrocytes featured lower overlaps, with the maximum number of shared terms being 13 between AD Ex 1 and PD Ast 1.

**Figure 4.**
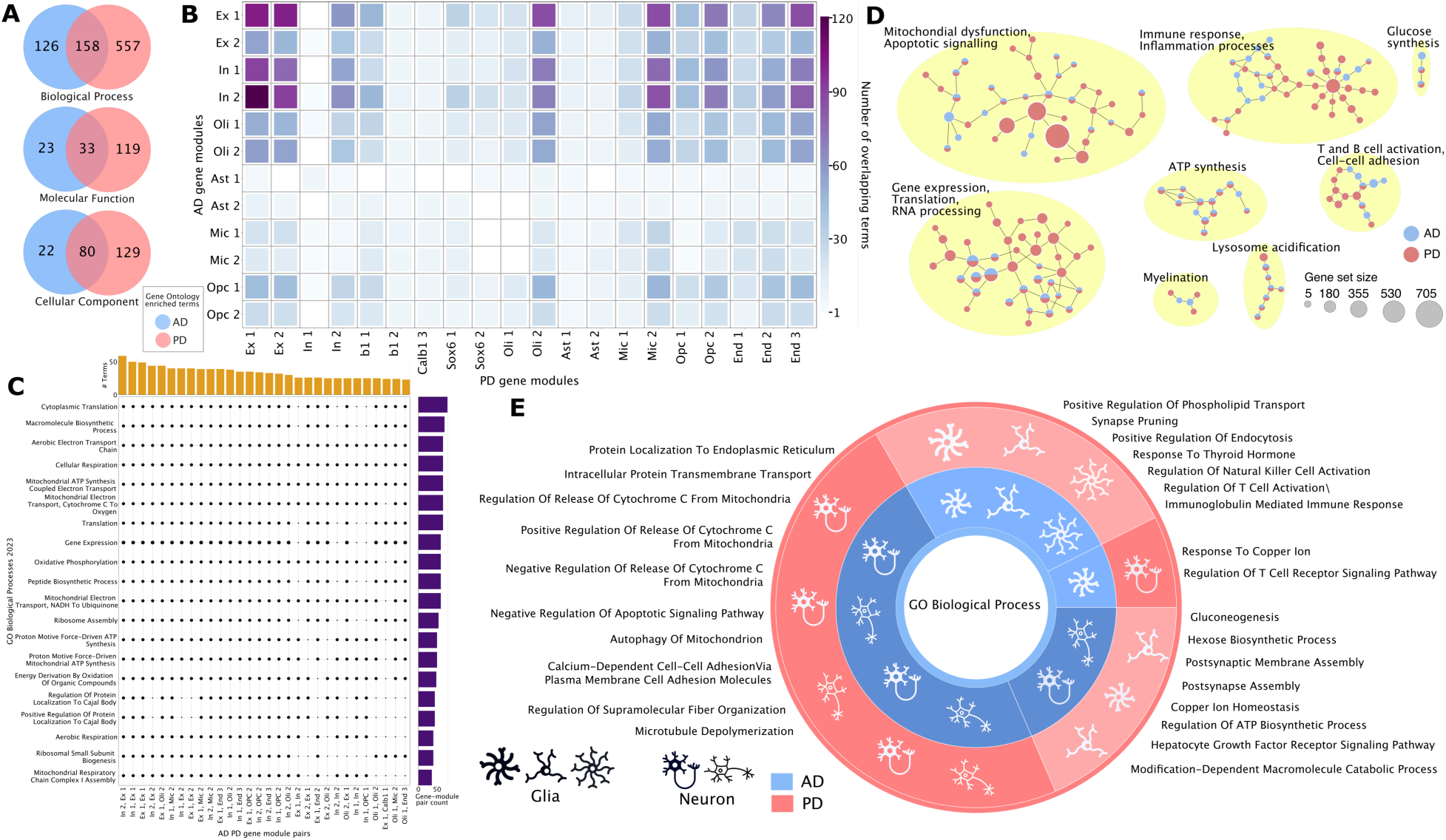
Gene ontology terms mapped to gene modules are shared between AD and PD. We performed gene set enrichment analysis on the gene modules derived from our previous analysis. For each gene module in AD or PD, we mapped the ranked genes (based on predictive weights) to terms in the gene ontology (GO) 2023 database. This helped ground the gene modules to pre-curated biologically relevant ontologies. **(A)** Overlapping terms from GO — Biological Process, Cellular Component, Molecular Functions. The venn diagrams depict the number of unique and shared terms across all gene modules, grouped by AD or PD. **(B)** Number of shared GO terms between every AD-PD gene module pair is shown. Brighter colors represent a higher number of shared terms. White grids represent zero overlapping terms. Neuron gene modules in both AD and PD had the highest number of shared terms, closely followed by oligodendrocyte-related module combinations from PD. **(C)** Zooming in on a few key terms summarizing the most frequent shared terms and gene-module combinations. Solid black dots indicate that a term (in the vertical axis) is enriched in the corresponding gene-module pair (in the horizontal axis). The bar plots on the horizontal axes are counts of the total number of terms common between AD and PD for the gene-module pair (arranged in decreasing order of term counts, first 30 pairs shown). Bar plots on the vertical axis represent the total number of cross-disease gene-module pairs that a term is present in (arranged in decreasing order of gene-module pair counts, first 10 terms shown). **(D)** Graph visualization of select biological processes across AD and PD from GO. Nodes are colored based on disease label and node size indicates the gene-set size. Groups names summarize the main themes from the terms in the group. This zoomed-out view highlighted key biological processes involved in both AD and PD. **(E)** Shared GO BP terms between AD and PD that are unique to broad cell type groups are shown. The inner circle denotes the cell type group from AD while the outer circle denotes the PD group. Darker shade represents terms enriched exclusively in neuron modules (excitatory and inhibitory neurons, CALB1, SOX6) and lighter shade represents terms enriched exclusively in glial modules (microglia, astrocyte, oligodendrocyte, OPC, endothelial cells). Biological processes related to altered cytoskeleton dynamics, impaired mitochondrial function, and apoptotic signaling are enriched across gene modules from neurons in both AD and PD. Immune response, synapse maintenance, and lipid transport-related terms are altered in one or more glial cell modules in both AD and PD.

By sorting all biological processes based on their frequency of shared occurrence across AD-PD module pairs, we identified the top 30 terms. These were related to protein translation, cellular respiration, and mitochondrial energy synthesis (Fig. 4C). To further summarize the terms systematically, we devised a visualization procedure to obtain a synoptic summary of the overarching biological processes. For this, we leveraged the predefined hierarchical tree structure of GO terms ^47^. Each node in the tree represented a GO term, and the edges described shared genes or functional relationships determined by careful experimentation. We subsetted this tree to 841 enriched GO BP terms derived across all AD and PD modules. Drawing these out in a network analysis tool, we extracted biological themes shared between AD and PD (Fig. 4D): protein synthesis and misfolding, immune response, lysosome acidification, glucose metabolism, mitochondrial dysfunction, and myelination.

For validation of these results, we turned to our external dataset pair. Analogously, we applied the GSEA pipeline to the gene modules derived from the Seattle-AD and Smajić-PD datasets. We successfully confirmed that AD and PD neurons, oligodendrocytes, and OPC modules had the largest occurrence of shared terms. Moreover, this also corroborated the broader biological themes of protein synthesis, mitochondrial energy metabolism, myelination, and glucose metabolism as shared between AD and PD (fig. S4).

Thus, our gene set enrichment analysis successfully identified a variety of matching biological, cellular, and molecular processes across AD and PD. We observed that the strongest interlocking effects emerged for gene module combinations that implicated neuronal cell populations, both inhibitory and excitatory, along with oligodendrocyte and OPCs. In contrast, we observed modest AD-PD overlap of terms from glial gene modules. Molecular signaling cascades related to stress, apoptosis, lipid metabolism, mitochondrial energy metabolism, and protein folding were among the top biological pathways that played a role in both AD and PD.

### Distinct glia and neuron-specific molecular pathways characterize AD and PD

In another overview analysis, we drilled down into the multitude of terms highlighted as being shared between AD and PD. We observed that the most frequently highlighted terms across all examined gene modules (cf. previous section; table S4) were shared between several cell types and were mainly cell injury response pathways. This finding aligns with existing knowledge that cellular responses to injury are often common across neurons, glia, vascular cells, and other systems of the central nervous system (CNS) ^48–52^.

To investigate the presence of cell type localized terms, we designed a probe to filter out potential cellular injury pathways. First, we binned the gene modules in our analysis into two groups – neuronal modules and glial modules. Neurons and glia are fundamentally different in terms of their anatomy and roles in brain function and thus are likely to exhibit distinct responses to disease. Our neuron set included gene modules from excitatory neurons, inhibitory neurons, CALB1, and SOX6, while the glia set comprised modules from astrocytes, microglia, oligodendrocytes, and OPCs. Next, we discarded all GO BP terms that were enriched in modules belonging to both neuronal and glial groups. Thus, the remaining terms had the property of being exclusive to either neurons or glia. We could then compare these terms and identified neuron and glia specific shared mechanisms between AD and PD (Fig. 4E).

Neurons shared the largest number of exclusive (unique to this category of cell types) GO BP terms between AD and PD. We identified several terms associated with microtubule depolymerization and cytoskeleton dynamics shared between PD and AD neuron modules (PD Calb1 1, Ex 1, Ex 2 and AD In 1, In 2 and Ex 1. Shared genes associated with these terms included MAPT, FKBP4, GBA2, MAP1A, MAP1B, MAP1S, MAP2, MAPRE3, STMN1, STMN2, STMN3, STMN4). Another highlighted biological theme involved alterations in the regulation of mitochondrial cytochrome c release. Ex 1 and Ex 2 in PD and In 1 and In 2 in AD highlighted these terms (PINK1, PRELID1, CLU, BNIP3, DNM1L, GHITM, GPX1, MFF, MLLT11, MOAP1). Alterations in iron homeostasis were noted in Ex 1 from PD and In 1 and Ex 1 from AD (SOD1, several ATP genes, CCDC115, FTH1, FTL, ISCU, NDFIP1, SLC22A17).

We noted that glia-exclusive terms had several themes centered around the immune and complement systems along with synapse pruning, lipid transport, metal ion homeostasis, and thyroid hormone balance. T cell activation was enriched in PD Mic 2, End 2, and AD Mic 2. These modules shared genes such as B2M, HLA-A, HLA-B, HLA-C, HLA-DPA1, HLA-DRA, HLA-DRB1, HLA-DRB5, and HLA-E. Lipid and phospholipid transport showed up exclusively in microglial modules, in AD Mic 1 and PD Mic 2 (APOE, TSPO, and PRELID1). Response to thyroid hormone was recorded in PD Mic 2 and AD Mic 1 (CTSB, CTSH).

We also noted a few terms exclusive to opposite categories in AD and PD. For example, “response to copper ion” was identified exclusively in AD glia modules and PD neuron modules. However, other functionally related terms like cellular response to copper ion, copper ion binding, and copper ion homeostasis, were enriched in AD Ast 1, Opc 1, Ex 1, and Ex 2 and in PD Mic 1, End 1, Ex 1, Ex 2, and In 1. A closer inspection of all terms belonging to cross-category modules, AD neuron-PD glia or AD glia-PD neuron, revealed that they were an artifact of GO’s granularity and naming convention. This was not the case for AD neuron-PD neuron and AD glia-PD glia specific terms. As an example, a manual search for “cytoskeleton” or “microtubule” highlighted only neuronal modules in both AD and PD. We also confirmed this finding using the GO BP results from Seattle-AD and Smajić-PD analysis arm.

Taken together, these observations highlighted key biological themes that appeared to be localized to certain cell types in AD and PD. Neurons in AD and PD demonstrated alterations to cytoskeleton structural integrity, mitochondrial transport, and mitochondrial energy synthesis. Alternatively, glia-based modules highlighted alterations to several support mechanisms including synapse pruning, lipid transport, and immune and inflammatory systems.

### The gene modules method outperforms traditional differential gene expression approach in linking AD and PD genetics

We compared our gene modules framework to the widely adopted univariate method in RNA-seq: DGE analysis. DGE identifies differences in gene expression between two groups (AD-control or PD-control in our case) by comparing expression profiles in diseased versus neurotypical cell states. We conducted a DGE analysis independently for the AD and PD datasets, focusing on each cell type separately (analogous to our main analysis). For each gene, we computed its log-fold change, contrasted between case and control, and identified a set of differentially expressed genes (DEGs). Statistically significant DEGs (FDR corrected p-value < 0.05; see Methods) were subject to further comparison between AD and PD cell types.

We estimated the overlap between the adDEGs and pdDEGs by computing Kendall’s tau-b (*τ*_b_) coefficient (Fig. 5) between the log-fold change values of significant DEGs for all combinations of cell types from AD (6 types) and PD (9 types). Among all the cell type pairings analyzed (6✕9 = 54 combinations), the highest correlation observed was 0.2, occurring between oligodendrocyte adDEGs and OPC pdDEGs. Notably, this maximum correlation among all possible pairings is significantly lower than the maximum *τ*_b_ observed in our gene module analysis (cf. Fig. 2A).

**Figure 5.**
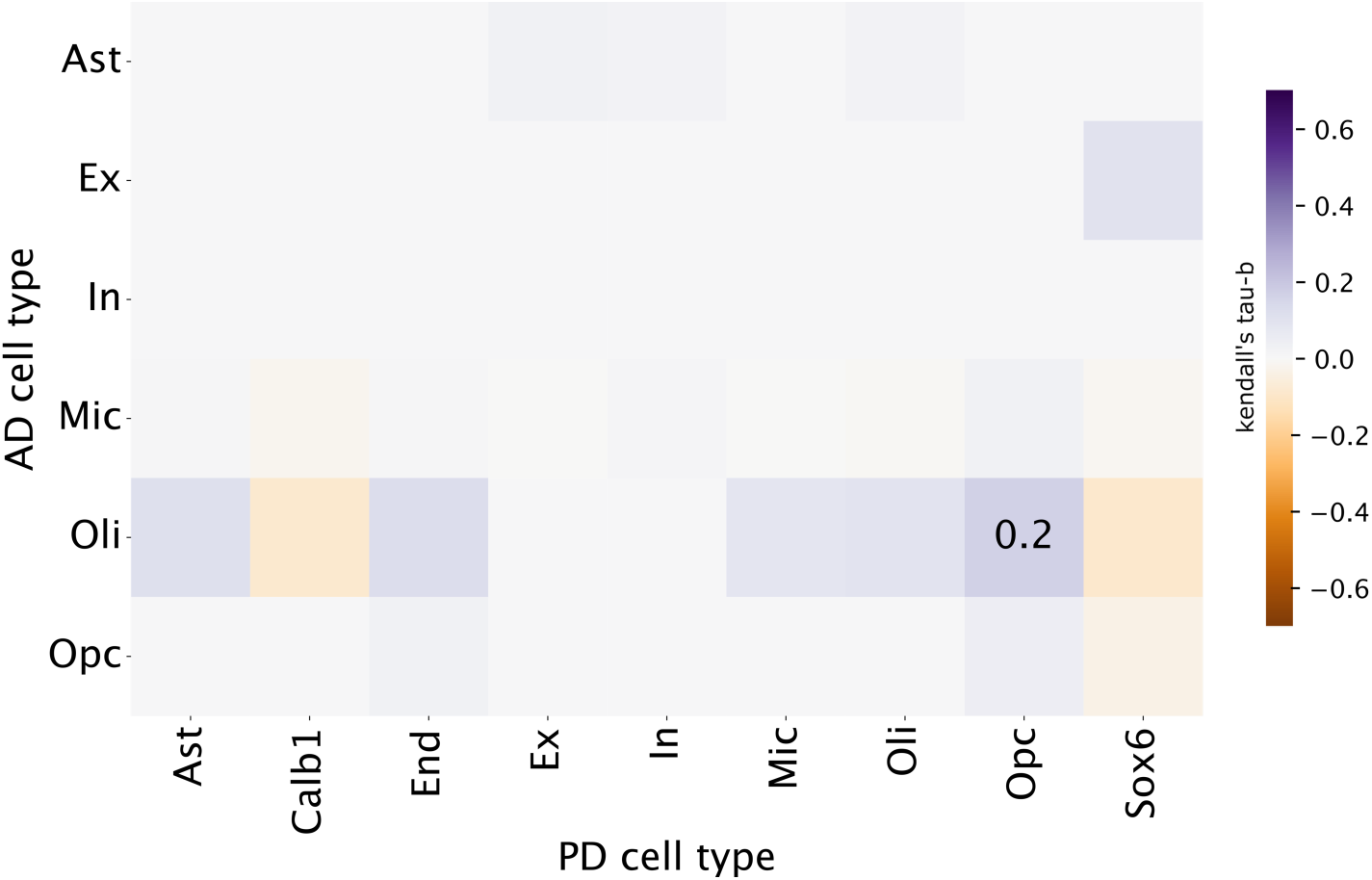
Legacy analysis approach, differential gene expression analysis, dulls in comparison to our gene module approach in identifying overlap in genetic mechanisms between AD and PD. To benchmark our latent factor modeling approach in identifying AD PD overlap, we conducted a differential gene expression analysis for AD and PD individually. The differentially expressed genes (DEGs) were identified using Wilcoxon rank-sum test contrasting transcription signatures between disease and control groups (adjusted p < 0.05 with Bonferroni correction for multiple testing; n=1000), conditional on cell types. Here, we compared the DEGs between AD and PD and depict their overlap between all cell type pairs. Darker colors denote higher Kendall’s tau-b of the log-fold changes in significant DEGs for each cell type, compared between AD and PD. The maximum correlation observed is 0.2 for AD oligodendrocyte and PD oligodendrocyte precursor cells. Compared to our PLS gene-module-based overlap analysis (Fig. 2A), DGE extract significantly smaller correlations. Ast, Astrocyte; Ex, Excitatory neuron; In, Inhibitory neuron; Mic, Microglia; Oli, Oligodendrocyte; Opc, Oligodendrocyte precursor cell; End, Endothelial.

We then systematically quantified the overall difference in mean correlations between cross-disease associations based on pairwise gene module *τ*_b_ from PLS (12 x 20) versus pairwise cell type *τ*_b_ from DGE (6 x 9). We used Welch’s t-test, suitable for samples of unequal sizes. Our results indicated a significant difference in mean correlation strengths. PLS_cell_ *τ*_b_s were, on average, 3.88 points higher than DGE *τ*_b_s (p value< 0.001). This finding suggested that inter-disease *τ*_b_ coefficients of AD-PD similarity were systematically stronger in our present gene module approach compared to the classical DGE approach.

To explore the biological relevance of the DGE-derived gene implications, we performed a comparable gene set enrichment analysis of the GO databases, GO BP, MF, and CC. Independently for each disease, we created our ranked gene list based on the log-transformed fold change (cell type conditional) and used GSEA to identify significantly enriched terms. We observed 20 common terms, in total, between AD and PD across all 3 GO databases (fig. S5). These terms were a significantly smaller subset of the common terms that were implicated through our gene-module analysis. Notably, these terms were among the most frequently occurring terms in the PLS analysis (Fig. 4C), which implied that DGE primarily identified the strongest disease signals from the raw gene expression matrices.

Thus, compared to the incumbent single-gene DGE approach, we demonstrated that our latent factor modeling achieved superior performance. This underscores the value of identifying cross-disease associations leveraging disease-relevant gene modules derived by analyzing the whole transcriptome in a single shot.

### GWAS-seeded co-expression networks also indicate AD-PD genetic overlap

In an alternative set of analyses, we pursued the same research question - the extent of AD-PD overlap - using a complementary quantitative analysis workflow. Devising a top-down framework (Fig. 6A) seeded with 164 GWAS genes (GWAS in AD or PD), we constructed disease-specific differential gene co-expression networks (DGCN) using the AD-Rosmap and PD-Kamath datasets.

**Figure 6.**
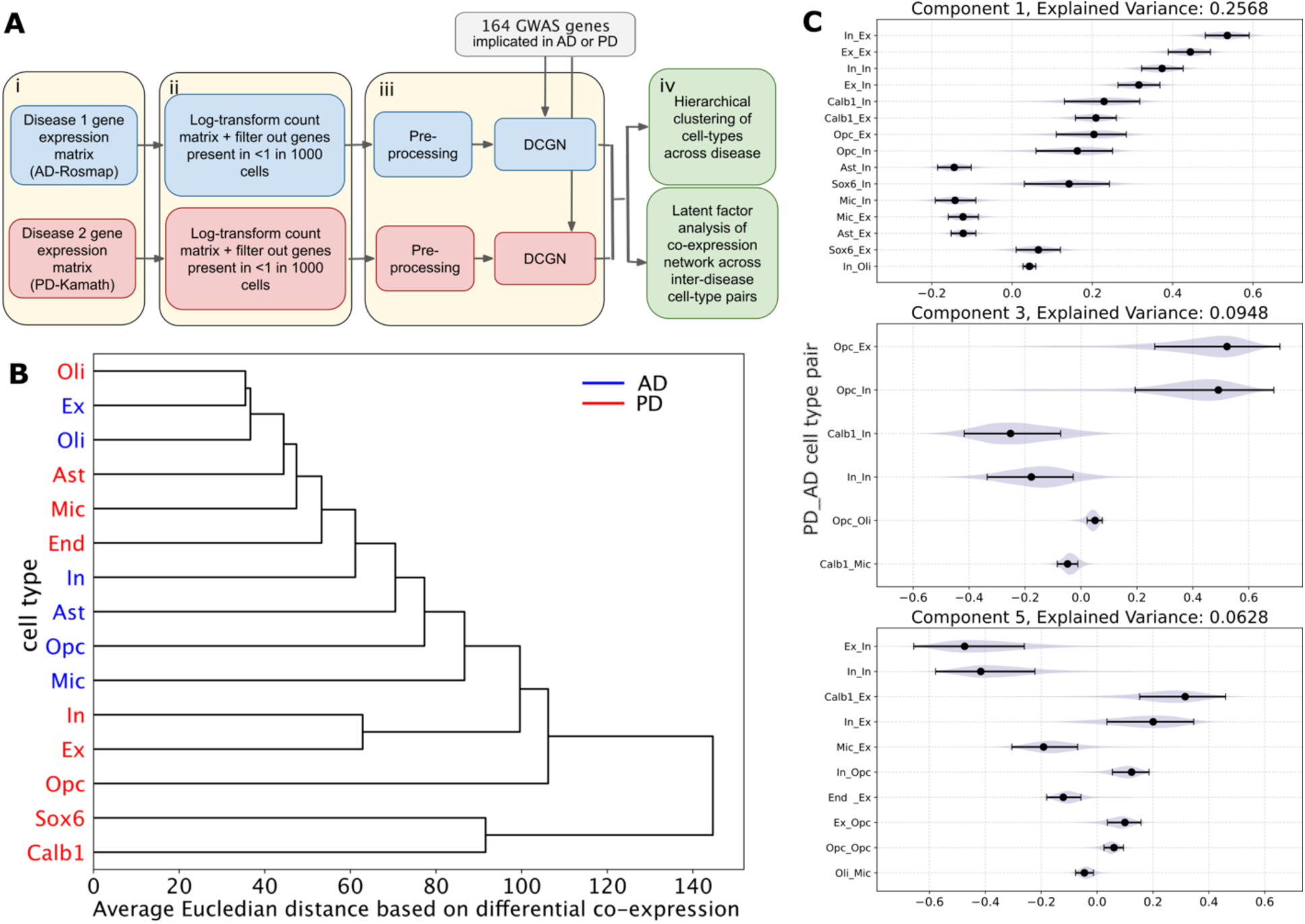
Differential gene co-expression network, a complementary analysis, identifies hints of overlap between AD and PD. To corroborate our findings in a technical replicate, we contrasted differential gene co-expression networks (DGCN) between AD and PD. We constructed DGCNs to identify genes (out of 16,936) whose expression patterns were harmonious with seed genes (164 GWAS genes from AD or PD), but whose synchrony was altered in disease compared to control conditions. (**A)** Overview of workflow. (i) Single-nucleus RNA seq datasets were downloaded from open-source repositories. Preprocessing done by source authors included quality control, normalization and cell type annotation. (ii) Local processing was performed independently for each cell type. This step followed recommended guidelines for data transformation and removed lowly expressed genes. (iii) Gene co-expression network analysis. For each gene from a set of GWAS implicated genes from AD and PD, a co-expression network was created for the disease and control groups independently. The results were subtracted to give the differential gene co-expression networks (DGCN). 164 seed genes, implicated by the largest AD or PD studies, were used to construct the DGCNs. (iv) Post DGCN analysis. The disease-specific DGCNs were compared between AD and PD to identify cross-disease cell type-cell type associations. **(B)** displays the hierarchical clustering of cell types based on the similarity of the co-expression patterns captured by DGCN. The dendrogram is based on the average Euclidean distance of the DGCNs for all cell type pairs between AD and PD. **(C)** Kendall’s tau-b (*τ*-DGCN) was used to calculate the pairwise associations between the DGCNs from two cell types. Cell type pair loadings for the top 3 principal components explain 50% of the total variance across all pairwise cross-disease DGCNs. Here, the pairs are presented in descending order based on the loading magnitude. The dots represent the empirical PCA loading. Error bars represent the 20/80% CI from 1000 iteration bootstrap analysis which sampled rows of a DGCN with replacement. Only pairs that did not include zero in their bootstrap interval are shown. The first and second principal components are dominated by cell type pairings involving mainly excitatory and inhibitory neurons in AD and PD. The second component emphasize PD dopaminergic neurons. The third component focusses on glial and vascular cell types from PD across cell types from AD. Ast, Astrocyte; Ex, Excitatory neuron; In, Inhibitory neuron; Mic, Microglia; Oli, Oligodendrocyte; Opc, Oligodendrocyte precursor cell; End, Endothelial.

The notion of gene co-expression networks (GCN) rests on the assumption that genes with similar expression profiles, across cell transcriptomes, often share functional or regulatory relationships ^53–55^. Our DGCN analysis integrated a contrastive element to GCN that compared disease and control groups. This allowed the elimination of the effects of housekeeping genes. Any residual patterns of covariation were then attributed to the effects of a disease. We computed DGCNs independently for the AD and PD datasets and separately for each cell type within a dataset. Concretely, for a given cell type, we computed Pearson’s correlation (*ρ*) between the expression profile for each GWAS gene and the expression signatures of the remaining genes (out of 16,936 transcripts), across all healthy cells, (matrix dimension 164 x 16,936). We then repeated the same analysis for the diseased cells (164 x 16,936). The difference gave us cell type-specific differential co-expression networks for each of the 164 GWAS genes for AD and PD separately.

To quantify the alignment between our two analysis arms, we measured the correspondence between the relevant genes from the DGCN analysis (top-down) and our PLS-derived gene modules (bottom-up). We looked at the number of robust genes that were common in each gene module - GWAS DGCN pair (fig. S6A). Our findings suggested high PLS_cell_ module-DGCN alignments for the same cell types. For example, co-expression signatures from astrocytes shared, on average (across 164 seed genes), 20% of genes (significant *ρ* at FDR < 0.01) with PLS_cell_ Ast 2. Similarly, excitatory neurons shared, on average, 58.4% of genes (significant *ρ* at FDR < 0.01) with PLS_cell_ Ex 1 and 42.2% of genes (significant *ρ* at FDR < 0.01) with Ex 2. The substantial module-DGCN alignment supports the intrinsic validity of both analysis approaches.

Finally, we proceeded to provide further elements to understand AD-PD overlap. We performed a hierarchical clustering of the combined and unraveled AD and PD DGCN matrix of dimension 15 (cell types) x 2,777,504 (seed-gene x transcriptome) (see Methods). The clustering grouped cell types from AD and PD based on their gene expression co-variation patterns centered on GWAS genes (Fig. 6B). Importantly, we did not combine the gene expression matrices; instead, the DGCNs were derived separately for AD and PD. As an overarching observation, our secondary analytical approach recapitulated the main observations from our primary latent factor modeling arm. Specifically, excitatory neurons from AD grouped closest to oligodendrocytes from PD, indicating that their underlying disease-related deviations in gene co-expression profiles were similar to each other. Oligodendrocytes also emerged as sharing similar co-expression profile alterations between AD and PD. We noted that microglia tended to be the furthest in the grouping within the respective disease. AD microglia showed the most distinct co-expression among other AD cell types, and similar observations held for PD. This recapitulated our primary analysis insight — microglia, from both AD and PD, were most singular in terms of the distinct functional pathways. Overall, the cross-disease similarity clustering noted here corroborated several key findings from our main analysis arm.

We then assessed the extent of AD and PD overlap in yet another way. We used Kendall’s tau-b to quantify the degree of similarity between the same seed-derived DGCNs from an AD-PD cell type pair (see Methods). Then we extracted a 2D embedding of the 164 (seed genes) x 54 (AD-PD cell type pairs) correlations using principal component analysis (PCA). The first 4 PCA components were determined to be statistically significant (0.5/99.5% CI; fig. S6B; see Methods). This examination elucidated any systematic cell pairings that tended to share similar co-expression profiles.

This derived embedding space highlighted interlocking cell pairs that exhibit overlapping co-expression patterns across different seed genes (Fig. 6C). The first principal component, accounting for 25.6% of the variance in seed-derived co-expression patterns, was dominated by robust cross-disease neuronal cell type pairs (subjected to bootstrap robustness check; see Methods). The second component (14.4% of total variance) grouped DA neurons in PD with excitatory and inhibitory neuron combinations from AD. This was followed closely by combinations of microglia in AD and oligodendrocyte and OPCs in PD in the later components. These results reflected similar conclusions as derived from our gene module-based findings (cf. above). Thus, our parallel analysis highlighted important patterns of overlap consistent with our PLS gene module-based findings, further validating our earlier results through a technical replication.

## Discussion

In this study, we examined the molecular, genetic, and cellular ties between Alzheimer’s and Parkinson’s disease. By analyzing the entire protein-coding transcriptome at single-cell resolution, our multivariate approach uncovered AD- and PD-deviant genes forming co-expressed cliques at sub-cell type granularity. We compared and quantified these gene modules demonstrating molecular overlap between AD and PD. Further, we mapped the gene modules to disease-relevant biological programs. This illuminated the complex genetic underpinnings leading to shared disease neuro-phenotypes. Thus, we provide single-cell genomics scientists with a tool to compare any pair of diseases from a global transcriptome perspective.

Our analytical protocol was enabled by access to an AD dataset (ROSMAP-AD) with 70,634 nuclei from 8 major cell types and a PD dataset (Kamath-PD) with 340,902 recorded nuclei from 11 major cell types. From both datasets, we extracted multiple gene modules for each cell type. These signified distinct modes of subcellular disease-deviated transcriptional changes. Our grading of the alignment of gene importance between pairwise gene modules from AD and PD demonstrated sizeable overlaps. In the spotlight were modules derived from AD or PD neurons, oligodendrocytes, and OPCs. This high degree of transcriptional differences mirrored between AD-PD was also highlighted in a secondary analysis arm, which compared gene co-expression networks between the two diseases. Additionally, we were able to replicate these findings in an external AD-PD snRNA-seq dataset pair. To our knowledge, these links are the first to reveal shared genetic control underlying measurable transcriptional differences in AD and PD.

To situate our findings within the context of GWAS, we examined whether established GWAS hits, from either AD or PD, were included in our gene modules. We found that these marker genes played robust roles within our functionally integrated gene modules despite being extracted in a transcriptome-wide approach. Additionally, we observed clear cell type localization of the popular genes within our modules. For example, APOE (the notorious AD risk gene) signatures were localized to microglia and astrocyte gene modules, agreeing with previous single-cell transcriptomics studies in AD ^30,37^. Central to our investigation, several AD-relevant marker genes surfaced in PD-associated gene modules and vice versa, highlighting the interconnectedness between the two disease categories. For example, APOE had robust PD associations in several of our PD gene modules, including those pulled from microglia, astrocytes, oligodendrocytes, and neurons. Indeed, APOE has previously been shown to be predictive of cognitive decline in PD patients based on clinical studies ^56,57^. Additionally, APOE has been shown to exacerbate PD pathology in mouse models ^58^. As another example, we consider SNCA, a major gene implicated in PD GWA studies ^59^. In the PD brain, misfolded SNCA protein, *α*-synuclein, is found in Lewy bodies and is a primary neuropathological marker ^60^. Here, along with several neuron and glial PD modules, SNCA emerges as a strong contributor to AD excitatory neuron modules. Indeed, APP transgenic mice with SNCA knockout have demonstrated a significant reduction in amyloid burden, hinting at connections between this PD gene and AD pathology ^61,62^. Taken together, we have successfully replicated the roles of important AD or PD genes while placing them within the fuller context of gene modules. We have further expanded the implication of these genes to cross-disease correspondences, to which previous univariate approaches were systematically blindfolded ^8^.

In addition to comparison with GWAS, we provided biological insights to our AD and PD comparison by grounding our gene modules in pathways from pre-curated gene ontologies. We identified shared alterations in cellular energy metabolism and stress response, inflammation, lipid signaling, protein folding, and protein degradation cascades. Some of these overarching disruptions in biological processes have been discussed in reviews summarizing the collective understanding from decades of neurodegeneration research ^48^. In a clean bottom-up workflow, our study confirmed these prior findings while going beyond previous studies by carefully localizing these molecular processes to cell-type-specific gene modules. For example, one potentially shared feature of neurodegeneration, as evidenced in previous brain tissue staining, was the presence of abnormal protein aggregates ^48,63^. These can vary in the type of protein (tau, amyloid-β, *α*-synuclein), the form of the aggregate (beta sheets, fibrils, oligomers), or the cell type they plague (neurons, astrocytes, oligodendrocytes). In line with this, our enrichment analysis revealed that protein misfolding (ER stress) and its associated biological processes were widespread across all major cell types, diseases, and datasets. We also detected extensive alterations in molecular pathways related to protein degradation (ubiquitin protein ligase binding and clathrin-mediated endocytosis ^64^), indicating a potential breakdown in protein disposal systems. Indeed, cellular assays had identified malfunctions in ubiquitin-dependent protein clearance in neurodegeneration ^65,66^.

### Shared molecular mechanisms between AD and PD in neurons

Neurons are particularly sensitive to proteasomal turnover due to their longevity and delicate synaptic regulatory requirements ^64^. The toxic effects of accumulated proteins ultimately lead to neuron death and usually mark the final stages of neurodegeneration. Zooming in on neurons, we identified precise mechanisms shared between AD and PD that were exclusive to neuron gene modules. These were primarily related to alterations in cytoskeleton dynamics, impaired mitochondrial functions, and apoptosis mechanisms.

Defects in cytoskeleton dynamics as major contributors to neuronal death have been cited by several lines of evidence, including microscopy and genetic studies ^67^. In our gene module enrichment, we found microtubule-associated processes to almost uniquely localize to neuronal modules in both AD and PD (in all 4 examined datasets). Alterations to post-translational modifications of neuronal microtubule’s acetylation levels have been reported before by in-vitro and in-vivo studies in both AD and PD ^68,69^. The involvement of the MAPT gene in these gene modules was noteworthy. This gene encodes for the protein tau which is responsible for stabilizing the axon microtubules. In cell culture studies, tau aggregates demonstrate prion-like behavior (prion-like characteristics of misfolded proteins imply the ability of self-propagation through seeding), passing from neuron to neuron across synapses ^70–72^. Taken together, the convergence of our findings with previous research hints at dedicated mechanisms that are possibly aligned between AD and PD and which can ultimately lead to prion-like manifestations ^73,74^.

Of note, the same gene modules that encoded dysfunctional cytoskeleton dynamics were also enriched in several terms related to the mitochondria, including axonal transport of mitochondria and protein localization to mitochondria (all 4 datasets). This observation alludes to a vicious cycle between dysregulated microtubules and impaired mitochondrial transport accompanied by oxidative stress. Intact microtubules in neurons serve as highways for cargo transport (proteins, mRNA, organelles like mitochondria) through axons and dendrites ^75^. Previous research have shown that alterations to its stability leads to the breakdown of mitochondrial transport, which in turn leads to mitochondrial reactive oxidative species (ROS) ^50,76^. ROS, in turn, exacerbates the levels of free tubulin ^77^ which in turn has been shown to interact with proteins like *α*-synuclein, promoting the formation of oligomeric aggregates in the form of Lewy bodies ^78^, or tau, promoting intraneuronal neurofibrillary tangles (NFTs) ^76^. Notably, in PD pathogenesis, *α*-synuclein oligomers are also highly effective at seeding further protein misfolding in a prion-like fashion ^79^.

Cells maintain a state far from thermal equilibrium by continuously extracting energy from their environment. Neurons, in particular, have especially high energy demands, and mitochondria are a major source of energy in these cells. We have observed alterations in the regulation of cytochrome c release and the apoptotic signaling pathway in several AD and PD neuronal gene modules. In the intact human cell, cytochrome c is present in the mitochondrial intermembranous space. The presence of oligomeric forms of amyloid-β, *α*-synuclein, and tau has been shown to increase mitochondrial membrane permeability. It can cause leakage of cytochrome c into the cell cytosol ^80^. The loss of cytochrome c from mitochondria compromises the downstream functionality of cytochrome c oxidase which is used for ATP energy production. Instead, once in the cytosol, cytochrome c is known to initiate mitochondria-mediated apoptosis ^81^. This form of neuronal death has long been suspected as an early event in the pathophysiological cascade leading up to both AD ^82^ and PD ^83,84^. Specific to PD, in a self-reinforcing manner, impaired cytochrome c release from mitochondria is thought to escalate *α*-synuclein oligomerization via radical formation ^79^. Together, impaired mitochondrial functions along with a disrupted axonal transport system may individually or collectively be a universal mechanism of neurodegeneration in AD and PD.

### Oligodendrocyte and Oligodendrocyte precursor cell modules in AD and PD

In oligodendrocytes and OPCs, we located additional gene modules with high AD-PD overlaps. Three key observations stand out related to these modules. First, we observed that a sizeable portion of GWAS genes localized to cross-disease oligodendrocyte modules. Notably, PD oligodendrocyte modules contained more GWAS genes than modules from other cell types. Previous transcriptomic studies have pointed out that several disease risk loci from GWAS are associated with oligodendrocytes in both AD ^85^ and PD ^86^. Our study confirms and expands on this observation. Second, in our enrichment analysis, several biological processes revolving around myelination and regulation of axonogenesis were specific to these cell type modules and were shared between AD and PD. Correlative macroscopic brain-imaging studies have linked deteriorating myelin health to AD progression ^87^. A recent invasive experiment posited a causal link between aging myelin and AD ^88^. In a few AD mouse models and a human PD model, RNA-seq analysis identified changes in oligodendrocyte transcription related to impaired myelination ^32,89,90^. Finally, here, in addition to strong intra-cell-type module associations, AD oligodendrocyte modules also exhibited a high degree of association with excitatory neuron modules from PD. Such similarity between oligodendrocyte and excitatory neuron transcriptional modifications in AD was previously reported in the ROSMAP-AD transcriptomic study, in terms of shared DEGs ^30^. The extension of this overlap to cross-cell-type modules from AD and PD suggests pervasive crosstalk between excitatory neurons and oligodendrocytes in neurodegeneration.

### Shared role of heavy metals is highlighted between AD and PD

In general, dysregulated homeostasis of heavy metals leads to an increased risk of the onset and progression of neurodegenerative diseases, as evidenced by studies in both humans and animals ^91,92^. Astrocytes, in particular, have been shown to remove excess heavy metals from the brain parenchyma ^93^. In our study, gene modules from astrocytes showed exclusive enrichment for response to zinc ions in both AD and PD. Several metallothionein (MT) genes were common between these modules (MT1E, MT1G, MT2A). Previous in vitro models have shown that zinc induces harmful A1-type reactive astrogliosis which promotes synaptic degeneration in neurons ^94^.

We found consistent enrichment of copper (Cu) ion regulation and interaction in both neuronal and glial modules across all examined AD and PD datasets. These modules featured several MT genes (MT1E, MT2A, MT3), and APP. In the brain, unregulated Cu readily derails redox chemistry (Fenton/Haber-Weiss reactions), thus forming more cytotoxic ROS ^95–97^. Biophysical and biochemical experiments underscore our metal pathway findings, showing that altered Cu ion levels triggered misfolding of *α*-synuclein ^98^ and amyloid-β ^99^. Interestingly, in our study, a distinct set of genes related to copper ion binding appeared exclusively in the excitatory neuron modules from AD and PD. These genes were linked to critical antioxidative enzymes (SOD1, PARK), SNCA, and the copper chaperone protein (ATOX1). Prior in-silico analyses of microarray data on brain tissue had reported a similar grouping of copper-handling genes (metallothionein group and the enzyme binding group) ^100^. Using our analytical framework, we were able to localize these effects to specific gene sets within distinct brain cell populations.

Another metal ion, iron ion, homeostasis terms were found to be enriched in neuron gene modules from both AD and PD. Across all datasets, excitatory and inhibitory neuron gene modules were involved. In the brain, iron plays a key role in myelin synthesis, neurotransmitter production, and overall metabolism ^101^. However, elevated levels of redox-active iron, often originating from degenerating mitochondria, accumulate in several neurodegenerative diseases as evidenced by biochemical, cell biological, and transgenic animal studies ^102–105^. Thus, understanding disruptions of these heavy metal pathways that extend across AD and PD is an interesting avenue for further investigation.

### Common mechanisms of microglia’s involvement in AD and PD

In contrast to neurons, overlapping microglia-implicated terms in AD and PD were highly diverse in their functional themes. This underscores the broad roles that these cells play in neurodegeneration, transcending brain regions. We posited that the difference in developmental origins of microglia compared to other CNS cell types, such as neurons and macroglia (astrocytes and oligodendrocytes), may explain a part of this phenomenon. Unlike neurons and macroglia, which derive from the neuroectoderm ^106^, microglia originate from peripheral mesodermal tissue ^107,108^ and exhibit considerable homogeneity across brain regions. Most microglial genes exhibit similar expression patterns extending across cortical regions (origin of our AD datasets) and striatal regions (origin of our PD datasets) ^109–111^. This uniformity in gene expression might explain why several biological functions altered in the degenerative state were shared between AD and PD in microglia modules.

Functionally, microglia are the primary immune cells of the CNS ^112^, and neuroinflammation and immune system dysfunction are believed to be key components of neurodegeneration ^113^. Consistently, microglial gene modules from our study were enriched for several immune-related processes. For example, we noted T cell activation in both AD and PD (all 4 datasets). Microglia, upon activation by neuronal stress, are thought to release pro-inflammatory cytokines and upregulate MHC class I and II molecules ^114^. Further, the inflammatory cytokines can induce the expression of adhesion molecules on brain endothelial cells, compromising the integrity of the blood-brain barrier (BBB). This BBB breakdown accelerates peripheral immune cell entry. Thus, in a chicken-egg scenario, microglia and endothelial cells drive a chain reaction of T cell activation, oxidative stress, and neuroinflammation ^115–117^. This domino effect may have been captured in a PD endothelial module which featured T cell activation simultaneously with biological processes like leukocyte adhesion to vascular cells, blood vessel morphogenesis, and diameter maintenance, pointing to the dysregulation of the BBB. This cascade of immune response events exacerbates ROS production and neuronal damage ^115,116^.

Another contributing factor to increased ROS is imbalances in thyroid hormones (TH) ^118^. We found a robust overlap for response to TH in microglial gene modules from both AD and PD. Epidemiological evidence have linked multiple thyroid-related autoimmune diseases to increased prevalence of both AD ^119,120^ and PD ^121^. However, the exact contribution of TH to either AD or PD pathophysiology has not been fully established. Recently, an AD mouse model has linked brain hypothyroidism with reduced microglial reactions to inflammatory stimuli and aberrant amyloid-β ^122^. Following this, an *α*-synuclein PD mouse model found connections between TH and glucocerebrosidase activity in microglia ^123^. Our findings that highlight TH involvement in AD and PD demonstrated the efficacy of our data-driven model in opening important avenues for investigation.

Our approach also mapped microglial gene modules from AD and PD to synapse pruning. These modules implicated key genes, including TREM2 and several C1q genes from the complement system, which is an integral component for synaptic refinement. Synapse loss is frequently observed as an early event in animal models of neurodegeneration ^124–126^. Variants in the TREM2 protein and aberrant activation of C1q genes have been shown to cause irregular synapse pruning in studies on AD mouse models ^126–128^. In a PD mouse model, suppressing TREM2 gene products in microglial cells was shown to accelerate the loss of DA neurons ^129,130^. Yet, the totality of glia-synapse interactions, especially the role of the complement system in PD, is under-investigated ^131,132^. Once again, our bottom-up approach identified a common disease outcome in AD and PD in the form of synapse loss – a potential area for further investigation.

In addition, lipid transport regulation was enriched in both AD and PD microglial gene modules (lipid, phospholipid, cholesterol, and sterol transfer terms) implicating key genes including APOE, TSPO, TMEM30A, several ATP-binding cassette subfamilies (ABC) genes (ABCG1, ABCA5) and NPC2. Excess lipids from dysfunctional and stressed neurons are transported to glial cells via apolipoproteins E and D ^133^. This excess lipid has been reported to accumulate as droplets in human iPSC-derived microglia, reducing their phagocytic capabilities and increasing the secretion of pro-inflammatory cytokines ^134,135^. To sum up, the above impairments mediated by microglia ultimately exacerbates ROS burden and neurotoxic buildups leading to neurodegeneration.

In conclusion, our proof-of-principle investigation of the transcriptomic terrain intersecting AD and PD identified and characterized key shared genetic components of neurodegeneration. We were able to (i) quantify the degree of overlap between unique gene modules from AD and PD, (ii) chart the extracted molecular changes in brain tissue to distinct cell types that matched in AD and PD, as well as (iii) interpret our in-silico derived gene module functional associations with several lines of existing experimental evidence. Future work can expand our study to include a greater diversity of neurodegenerative, neurodevelopmental, and psychiatric diseases. Moreover, applying a similar framework to different, readily accessible source transcriptomes, like blood or cerebrospinal fluid, can identify critical biomarkers for neurodegeneration.

## Materials and Methods

### Rationale

Alzheimer’s disease (AD) and Parkinson’s disease (PD) have traditionally been studied as distinct entities, largely due to differences in their etiological triggers, neuropathology, and their association with specific brain regions. AD typically follows a progression that begins in the neocortex ^13,136,137^, while PD is often associated with dopaminergic neuron death in the substantia nigra ^31^. However, recent research has pointed to potential molecular and cellular overlap between the two diseases, suggesting the involvement of evolutionarily conserved pathways that operate across different cell types in the central nervous system ^48,51^. These shared mechanisms may give rise to similar neurodegenerative processes despite the distinct clinical presentations of AD and PD.

Existing approaches, such as GWAS, DGE, and other univariate regression analyses, have focused on identifying single-gene effects in isolation. While informative, such methods are inherently limited, as they do not fully capture the complexity of gene regulation, which occurs within tightly regulated networks of co-expressed genes. This is especially true in complex diseases like AD and PD, where risk genes may not always overlap. However, similar pathways of disease progression, involving neuron death and immune responses might be shared.

Given the modular nature of gene regulation, where genes work in concert within networks ^22^, we argue that studying gene expression through a multivariate lens (as “cliques”) offers a more accurate understanding of the molecular mechanisms underlying AD and PD. We propose an approach that extracts disease-relevant gene modules from single-nucleus RNA sequencing (snRNA-seq) data, which provides cell-type-specific insights at high resolution. By capturing the combinatorial nature of gene regulation, we aim to uncover potential co-occurring molecular cascades within cells that could be driving both AD and PD pathology.

This study tested two core hypotheses: (i) gene co-expression cliques, rather than individual genes, might offer a more comprehensive framework for understanding the genetic overlap between AD and PD, and (ii) within each cell type, multiple molecular pathways may contribute to disease, given the combinatorial nature of gene regulation. Our approach leverages supervised latent factor modeling, which has previously demonstrated effectiveness in identifying gene signatures in AD ^35^. Here, we apply this method in a comparative analysis of AD and PD to explore the molecular commonalities and differences between the two diseases at a transcriptomic level.

### Single genomics data resources

The advent of single-cell RNA sequencing has revolutionized cell biology by enabling the definition of cellular identity and heterogeneity through transcriptome data in both healthy and diseased tissues ^138–140^. As a variant of this, single-nucleus RNA sequencing (snRNA-seq) is particularly well-suited for tissues like the brain, where the availability of fresh tissue samples is limited ^141,142^. Our present study benefited from recently emerged and expansive snRNA-seq data, pertinent to AD or PD, from four separate and independent studies.

### Primary datasets

ROSMAP Alzheimer’s dataset ^30^, GEO accession number **<u>GSE178265</u>**: We examined an exceptionally valuable dataset of gene expression – the first investigation of AD using snRNA-seq. This dataset was derived from post-mortem brain tissue sourced from the prefrontal cortex (BA10) of individuals participating in the Religious Orders Study or the Rush Memory and Aging Project (ROSMAP) ^38^. All participants enroll without known dementia and agree to annual clinical evaluation and brain donation. Both studies were reviewed by an Institutional Review Board of Rush University Medical Center, and all participants signed informed and repository consents and an Anatomic Gift Act. The dataset was collected from 48 subjects who were carefully matched in terms of age and sex. These subjects consisted of 24 males and 24 females, with an equal amount of 24 individuals diagnosed with AD and 24 control subjects. The mean age of the individuals was 85 years. The recorded cell types included inhibitory neurons, excitatory neurons, oligodendrocytes, oligodendrocyte precursor cells, astrocytes, microglia, endothelial, and pericytes. The dataset comprised a total of 70,634 cell transcriptomes from 8 cell types and encompassed transcript counts for 17,926 protein-coding genes, aligned with the human reference transcriptome (hg38 GRCh38.p5).

Parkinson’s disease, Kamath dataset ^33^, GEO accession number **<u>GSE178265</u>**: This dataset included snRNA transcriptomes from post-mortem human midbrain and dorsal striatum (caudate nucleus and substantia nigra) tissue. Tissue samples were derived from an age and sex-matched cohort of 15 subjects, consisting of 7 males and 8 females. Among this group, 6 individuals had been diagnosed with PD, while the remaining 9 were controls. The mean age of the individuals was 83 years. Major cell types in this dataset included inhibitory and excitatory neurons, oligodendrocytes, oligodendrocyte precursor cells, astrocytes, microglia, and endothelial cells. The dataset also recorded a specialized subset of dopaminergic neurons from the midbrain, CALB1, and SOX6. The dataset provided transcript counts for 33,692 genes (genes were aligned to hg19) from 11 cell types and encompassed 340,902 single nuclei transcriptomes.

### External validation datasets

Seattle Alzheimer’s dataset ^37^: The authors of this resource sourced brain specimens from the Adult Changes in Thought Study and the University of Washington’s Alzheimer’s Disease Research Center. It is worth noting that the cohort detailed in this research represents the entire spectrum of Alzheimer’s disease severity. Brain tissue samples were drawn from the middle temporal gyrus. The study included participants from age groups ranging from less than 65 years to more than 90 years. To maintain consistency with our other datasets, we considered only participants over the 65 to 77-year age bracket. This gave us 76 individuals, with 47 females and 29 males. Out of these, 36 subjects were cases of recorded dementia, and 40 were controls. The mean age of the individuals was 88 years. The preprocessed datasets for the cell types labeled ‘L4 IT’, ‘L5 IT’, ‘Vip’, ‘Pvalb’, ‘Sst’, ‘Sncg’, ‘Oligodendrocyte’, ‘Microglia’, ‘Astrocyte’ and ‘OPC’ were downloaded from the <u>cellxgene platform</u>. The final dataset featured 683,260 nuclei across 10 cell types and recorded 36,517 genes.

Parkinson’s disease, Smajić dataset ^32^, GEO accession number **<u>GSE157783</u>**: The creators of this resource worked with post-mortem midbrain tissue sections and linked with the associated clinical and neuropathological data from the Parkinson’s UK Brain Bank and the Newcastle Brain Tissue Resource. The dataset includes age and sex-matched nuclei samples from adult human post-mortem midbrain tissue from 5 cases of idiopathic Parkinson’s disease, all of whom exhibited severe neuronal loss in the substantia nigra and had no family history of the disease. 6 control midbrain tissue samples were also collected to match the characteristics of the idiopathic Parkinson’s disease patients. The mean age of the subjects is 80 years. In total, 24,005 genes (mapped to hg38) were recorded in 41,435 single nuclei from 10 cell types.

### Preprocessing pipeline at source

We relied on the preprocessed datasets from the authors responsible for the data collection (cf. above). This maximizes reproducibility and compatibility with other studies working with these resources. The transcriptomic datasets were processed in the source studies using standard snRNA-seq processing pipelines. This included quality control for cell inclusion, including doublet detection, the removal of low-quality and outlier cells, the removal of lowly expressed genes, and cell clustering into cell types (exact details can be found in the Methods sections of individual research). Importantly, the cell type classification - which cell belongs to which cell population - was taken from the original studies as a basis for our investigations.

### Primary local preprocessing

To maintain comparability across the different datasets that recorded a variable number of genes, in our study we looked at the effects of genes that were recorded in all 4 AD and PD datasets. This gave us a total of 16,936 protein-coding genes that we used in further analyses, enabling us to make an apples-to-apples comparison. By considering only protein-coding transcripts, we also reduced our feature space size which is beneficial for any high dimensional analysis ^143,144^.

### Identifying gene modules: supervised latent factor modeling

At the heart of this study, we wanted to explore synchronous gene expression changes that occur in the brain accompanying a disease state and compare them between AD and PD. Supervised latent factor models are a natural choice to uncover the hidden patterns in a high-dimensional feature space, simultaneously accounting for the relationships between the observed variables and their associated disease-vs-control labels. Here, gene transcription signatures were the input variables that are to be embedded in a space maximizing disease versus control class separation. Gene expression measurements are known to have high correlations among themselves ^21^, giving rise to an additional modeling consideration (multicollinearity). Partial Least Squares Regression (PLS-R) ^145^ is a multivariate statistical technique that is particularly well suited in situations with multicollinearity (high correlation among features) or when dealing with high-dimensional data with considerable noise and small sample size.

In the present analysis, we utilized a variant of PLS-R for classification with one dependent categorical variable. This is known as PLS discriminant analysis (PLS-DA) ^146^. In doing so, we developed a predictive modeling framework to categorize disease versus control diagnosis based on input gene expressions. Formally, our input datasets are denoted by, 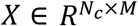 and 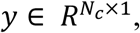 where *N*_*c*_ was the number of observations (nuclei) for a cell type *c*, *M* the number of measured features (genes; *M* = 16,936), *X*_*i*,*j*_ holds the recorded transcript count for the *i*^*th*^ nuclei and *j*^*th*^ gene, and *y*_*i*_ was the disease label (+1 for disease and -1 for control). The goal was to construct a low-rank projection from the 16,936-dimensional gene space that maximized the within-class separation between disease and control.

Concretely, PLS-DA can be viewed to consist of two key equations:

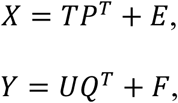

where *T* and *U* are *N*_*c*_ × *k*_*c*_ score matrices of *k_c_* extracted components, *P* is a *M* × *k*_*c*_ (gene-wise) loading matrix (effect size) of *X* and *Q* is a 1 × *k*_*c*_ loading vector of *Y* respectively. *E* and *F* are the residual matrices of *X* and *Y* respectively. The decomposition of *X* and *Y* is set to the solution of the optimization objective:

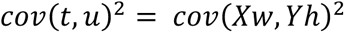

where *cov*(*t*, *u*) is the captured covariance 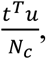 *w* and *h* are weight vectors that are extracted using the NIPALS algorithm ^147^. Interpreting loading vectors in PLS classification is essential to understanding gene expression features’ impact on disease detection. High absolute loadings signal strong contributions (positive to the target disease, negative to the control group), while near-zero loadings indicate minimal impact ^35^.

PLS-DA is prone to data overfitting when the number of features far exceeds the number of observations, especially in the case of inherently noisy, highly collinear single-nucleus data ^148^. To address this issue, we first applied principal component analysis (PCA) for dimensionality reduction to the gene expression input matrices. Applying dimensionality reduction techniques has been merited as a pre-processing for PLS-DA ^149^. The optimal number of PCA components was kept to *min*(500, *N*_*c*_), where *N*_*c*_ was the number of recorded nuclei for the given cell type *c* in the dataset. We used this transformed gene space for the PLS-DA analysis described above. To transition from the PCA embedding space back to the gene space, we project the PLS loadings (in the lower dimensional space) back into the original gene expression space. This ensured that the domain interpretability of the PLS estimates was preserved in the biological ambient space.

### Model selection, training, and performance assessment

Different cell types perform widely diverse functions, each arising from the functional recruitment of distinct gene families. In our pursuit to discover biologically meaningful gene groups, we implemented our quantitative analysis pipeline for a given disease on a cell type-by-cell type basis. This agenda enabled us to extract coherent latent components (latent factor/loading vector, hereby referred to as gene module) specific to each cell type. To ensure an even sample size in the disease and control group, we randomly sub-sampled transcriptomes of a given cell type to have comparable counts of nuclei from both disease and control instances. On this sub-sampled data, we removed genes that were captured in fewer than 1 out of 1000 cells to reduce our model’s degrees of freedom (*scanpy.pp.filter_genes*), which is in line with previous research ^150,151^. To reduce technical variation from sequencing depth, we normalized the data by dividing the raw UMI count by the total number of detected UMIs in each cell (*scanpy.pp.normalize_total(*data, target_sum=1e4)). Next, to account for the heteroskedasticity originating from differences in highly expressed vs lowly expressed genes, we log-transformed the normalized data from the previous step (*scanpy.pp.log1p*(data)). These dataset normalization approaches have been shown to work well as a preparatory step for downstream dimensionality reduction ^152^. In a data cleaning step, inter-individual variation in gene expression that could be explained by differences in postmortem interval was regressed out ^153,154^. The thus derived balanced, cleaned, and standardized transcriptomic profiles for each examined cell type were used for subsequent steps in our pattern-learning analysis pipeline.

After preprocessing and cleaning the transcriptomic data resources, we carried out the estimation or training of our supervised learning model. The PLS classifier needs tuning of one hyperparameter - the number of hidden components of variation in the data. Selecting this optimal number of latent components is crucial, choosing too few components implies losing out on crucial information and too many invariably leads to overfitting. We adopted a rigorous 10-fold cross-validation (CV) scheme to inform this model selection problem, carried out separately in each cell type. The set of transcriptomes was randomly split into 10 equal-sized data point subsets. We ensured that the disease-to-control ratio of cells for each subset reflected that of the full dataset. Screening a range of component choices (1 to 8), in each iteration, 9 out of these 10 data subsets were combined and used for training a PLS model, while the held-out subset was used for evaluating the choice of component number. The model’s performance was evaluated based on the area under the receiver operating characteristic curve (AUROC) in disease discrimination. This was performed for all combinations of training and validation subsets (*scikit-learn model_selection.GridSearchCV* function with *PLSRegression* as the ‘estimator’, ‘scoring’ set to ‘roc_auc’ and ‘n_components’ parameter set to 1-8). The number of components yielding the maximum mean AUROC over the CV subsets was noted as optimal for the given cell type. Note that this approach allowed us to independently derive the number of components that maximized disease-vs-control identification for each cell type, without overfitting to the input transcript space. This gave us 12 gene modules in AD across 6 cell types (2 cell types in ROSMAP-AD, pericytes and ependymal cells, did not pass our significance test. They were removed from further analysis) and 20 gene modules from 9 PD cell types (2 cell types in Kamath-PD, macrophages and ependymal cells, did not pass our significance test. They were removed from further analysis).

Next, we fitted PLS models on the full set of transcriptome observations for each cell type (PLS *model* specified using *sklearn.cross_decomposition* module *PLSRegression*). In doing so, we extracted multiple, unique disease-relevant gene modules. The statistical significance of an overall gene module was assessed in a principled, non-parametric permutation procedure. In 1000 permutation iterations, the transcriptome signatures were held constant, while the disease labels (outcome of model) were shuffled randomly across transcriptomes. The resulting surrogate datasets preserved the statistical structure of gene expression profiles while selectively destroying the association of the transcription profiles (model input) with diagnosis (model output). This approach generated a null distribution with minimal modeling assumptions ^155,156^. The empirical covariance (test statistic) between the gene expression and disease signature captured by each module (*cov*(*t*, *u*) defined above, *t* being *model*.x_scores_ and *u* being *model*.y_scores_) was compared with the resulting permutation distribution. This distribution reflected the null hypothesis of random association between gene transcription and the disease designation, which we test the actual model instance against. We deemed significant a module’s input-output covariance in the latent space if fewer than 5% of the null models yielded a better covariance strength than the original covariance from the actual model instance (fig. S1 C and D). In case a module failed to pass this label-shuffling permutation test, it was dropped from further analysis, along with all underlying gene modules for that cell type. Thus, in a data-driven approach, we were able to determine which gene modules in a cell type at hand carried enough information that allowed us to discern a biological signal from noise. Consequently, modules from cell types with few recorded nuclei were discarded, as their corresponding PLS models did not robustly pass the described rigorous permutation scheme.

To identify the subset of the examined genes that robustly contributed to disease detection in each gene module from a cell type, we implemented a 500-iteration bootstrap (BS) scheme. The bootstrap resampling was done by selecting nuclei, with replacement, from the cell type observations before applying dimensionality reduction. This approach simulated random nuclei sample draws that could have been derived from the broader cell population. Dimensionality reduction (PCA) and PLS model estimation were performed on the resampled bootstrap dataset in an identical fashion (cf. above). We disregarded any genes whose model coefficient effect size (PLS loading corresponding to this gene) included zero in its 5/95% BS-confidence interval (CI) for that module (i.e., its effect was removed by setting the loading value to zero). The derived ‘robust’ gene modules will henceforth be denoted as *P*^∗^(dimension *p* × *k*_*c*_; cf. PLS definition section above) in future references.

An inherent ambiguity of the class of latent factor models (i.e., aspects of model non-identifiability), including PCA and PLS, is the reflection invariance of derived latent vectors. To remedy this source of indeterminacy, we computed the cosine similarity (*γ*; range -1 to +1), between a BS loading vector and its corresponding empirical loading vector. In the scenario where *γ* was less than 0, indicating a flipped (“mirrored”) loading vector, we multiplied the loading vector of the BS model elementwise by -1 to align it with the original loading vector. This method has been employed by previous authors to address the issue of reflection ^35^. The resulting distribution of loadings for a gene in a module was compared to its counterpart in the original model estimate.

Motivated by our overarching goal to quantify the association between gene modules from two diseases, we first sought to explore the degree of similarity and difference between pairs of gene modules obtained from the same disease. Each gene module from a cell type captures a unique and complementary view of the relative roles of genes for successful disease prediction, hence we expected to see a low degree of association. In contrast, we did not rule out the existence of cross-cell-type gene module similarities. Kendall’s tau-b correlation metric (*τ*_*b*_; ±1 being a high degree of association and 0 being no association) was chosen to gauge this overlap signal (cf. rationale of *τ*_*b*_ metric discussed later). We calculated *τ*_*b*_ for all combinations of module-module pairs (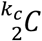 combinations, for *k*_*c*_ modules), resulting in a 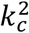 correlation matrix. A low magnitude *τ*_*b*_ indicated complementary modules whereas a magnitude of *τ*_*b*_ close to 1 indicated a higher degree of similarity in the compared gene cliques.

To audit the performance of the individual PLS models calibrated for each cell type in classifying disease versus control, we employed AUROC of disease classification as our evaluation metric. Given that cell samples from a patient can exhibit significant autocorrelation, it is crucial to account for this when evaluating the model. Traditional test-train splitting methods often involve blindly partitioning the dataset. This can result in overly optimistic test performance and makes it challenging to detect overfitting during the testing phase ^157^. In light of this, we employed a variation of cross-validation combined with bootstrapped Latin partitions ^158^ to ensure patient-level stratification. Concretely, in each iteration, a random sample of subjects (not cells) was drawn with replacement. This formed the basis of the training dataset’s nuclei source. Based on the subset of patients, a random sample of cells was drawn, with replacement, while ensuring that the disease-to-control ratio of nuclei was reflective of the empirical dataset. The percentage of cell samples from any given subject was also preserved in each iteration. This analytical protocol ensured that transcription signatures from the same patient were not present in the training and testing set at the same time. Individual PLS models were fitted on the re-sampled train dataset. This fitted model was then evaluated based on AUROC scores on the out-of-bag nuclei (test set), i.e., the transcripts from subjects that were not included in the training step. We performed 1000 iterations of this bootstrap-based model disease classification performance evaluation. This allowed for a principled assessment of the disease discrimination strength of the PLS solutions based on the cell type-specific gene transcription signatures.

### PHATE visualization

To gain a synopsis of the uniqueness of different gene modules from a single cell type, we created a concise, low-dimensional representation of high-dimensional cellular transcriptomes. For this purpose, we applied an especially flexible dimensionality reduction using PHATE ^39^. Compared to prevalent visualization techniques like tSNE or UMAP, PHATE is well suited for noisy snRNA-seq data. It has been shown to preserve both local and global structures in a dataset and can capture non-linear relationships in the transcriptomic information, by mapping them into diffusion maps relative to each other. We estimated a separate PHATE model for each cell type. We projected the transcriptomes from each cell type into independent low-rank spaces (using the *scanpy external.tl.phate* function with default parameters except n_pca = 500). Instead of using specific disease marker genes as done traditionally to identify subtypes for cells, we colored the cells in the PHATE embedding space based on their PLS score for each latent component (*PLSRegression x_scores_*).

### Quantifying disease overlap at the gene-module level

To quantify the similarity between AD and PD at the level of gene expression patterns, we computed the association between functional gene programs of AD and PD. Importantly, a similarity metric (correlation) was computed across the derived model’s predictive rules for a disease. That is, we did not pit the raw gene expression measurements against each other.

Formally, for a disease *d* and cell type *c*, the *i*th robust gene module (cf. above) can be denoted as 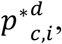 where *p*^∗^ is a vector of dimension 1 ⨉ *M* (number of genes in the input feature space). A robust gene module can contain tens to thousands of genes with non-zero loadings (out of ∼17,000 genes) while the remaining loadings are zero (cf. above). To minimize the impact of tied zero loadings in the correlation metric calculation between two modules, we considered only those genes that had non-zero weights in both modules (AND conjunction). This approach works well to identify groups of genes with similar disease contributions between two modules while ignoring genes that might have robust effect sizes in one disease but not the other.

Concretely, we employed Kendall’s tau-b ranked correlation metric (*τ*_*b*_) to evaluate pairwise correlations between AD and PD gene modules. Kendall’s tau-b correlation effectively handles tied ranks and provides a more accurate measure of ordinal association between gene modules. This is unlike Pearson’s-r, which assumes monotonicity and is unstable, or Spearman’s-r, which is biased and difficult to interpret ^159^. For all pairwise gene modules from AD (12 modules across all cell types) and PD (20 modules across all cell types), Kendall’s tau-b rank correlation coefficient, *τ*_*b*_O(*c*1, *i*), (*c*2, *j*)Q, was calculated using the scipy function *stats.kendalltau* 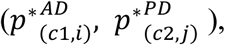 where (*c*1, *i*) is the *i*th gene module for the cell type *c*1 present in the AD dataset, (*c*2, *j*) is the *j*th gene module from the cell type *c*2 present in the PD dataset. To independently assess the statistical significance of each coupled module association, we employed a non-parametric permutation procedure with the null hypothesis of random association between gene modules from different diseases. For each gene module pair, we re-utilized the label shuffling derived module weights (cf. above) to calculate a null distribution from 1000 permutation iterations. We only interpreted a module-pair’s correlation coefficient that emerged as statistically relevant against a 5/95% CI threshold.

Yet another test was conducted to estimate the statistical sensitivity of the coupled associations of cross-disease gene modules. Across 500 iterations, we randomly bisected the empirical AD and PD datasets (before cleaning and standardizing) ensuring to preserve the original proportion of cell type nuclei and the disease-control ratio for each cell type, into two pairs of AD-PD subsets. Since the bisection reduced the effective observation sample size of each dataset to half, the downstream PLS fit enabled a suitable robustness check of derived gene modules. For each AD-PD subset pair, we ran our workflow pipeline (Fig. 1), steps A-D in parallel, resulting in 2 analogous sets of gene modules (a couple of 12 AD and 20 PD modules). We then performed Kendall’s tau-b correlation across these modules, giving us 2 correlation matrices of dimension 12 x 20. We unraveled these matrices and calculated Pearson correlated (*ρ*) of the absolute *τ*_*b*_ values. We used absolute values since we were interested only in association strength, not direction. In doing so, we essentially compared the *τ*_*b*_ correlation levels between different gene module pairs derived from complementary subset pairs. The estimated mean *ρ* gave us an indicator for the reliability of the quantified correlations between gene module pairs. A high *ρ* would signify a substantial degree of agreement between the association strengths of cross-disease gene modules, which in turn would attest to the robustness of the similarity analysis.

### Differential gene expression

Differential Gene Expression is ubiquitously used in snRNA-seq analysis to identify genes that show statistically significant differences in expression levels between two conditions or groups ^160^. It is a univariate method, meaning it examines each gene individually, thus losing key information hidden in gene co-expression patterns. We employed this traditional method to serve as an acid test for our new approach. Considering one gene at a time, we estimated the log-fold change of gene expression between the disease and control group ^161^. Log fold change is calculated by taking the logarithm (usually base 2) of the ratio of the mean expression levels of a gene between two conditions. To determine the statistical significance of these changes, we used the nonparametric Wilcoxon’s rank sum test, to help identify genes that had significantly different distributions in cases when compared to controls. We corrected for multiple testing using the Benjamini-Hochberg method ^162^. This gave us a set of statistically significant differentially expressed genes (DEGs; FDR<0.05). We did this separately for every identified cell type within a snRNA-seq dataset (*scanpy* tool *tl.rank_genes_groups* with method= ‘wilcoxon’). The final DEGs are referred to as adDEGs for AD and pdDEGs for PD.

To estimate the pairwise association between cell-type-specific adDEGs and pdDEGs, we employed Kendall’s tau-b correlation of the log-fold change values on the AND conjunction genes (cf. above). In doing so, we calculated a correlation matrix of dimension 6 (number of AD cell types) x 9 (number of PD cell types). To test for statistical significance, we performed a 1000-iteration permutation test. We randomized the disease label for each individual and subsequently identified a different set of adDGEs and pdDGEs. Based on the null hypothesis of a random association between AD deviant gene expression and PD deviant gene expression, we identified cell type pairs whose empirical association signal was significantly different in at least 95% of the 1000 permutation iterations.

### Quantifying difference in association strengths between PLS and DGE derived conclusions: Welch’s t-test

To formally quantify the cross-disease association information extracted by our latent factor approach versus DGE, we turned to a statistical test that can compare the central tendencies of the respective correlation measures. Welch’s t-test was a natural choice of method here as the number of correlated combinations being compared were different (240 *τ*_*b*_ gene module combinations from PLS and 54 *τ*_*b*_ cell type combinations from DGE), and the variances were not assumed to be equal ^163,164^. Welch’s t-test can be formally computed as:

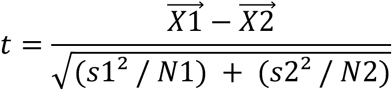

In our case, 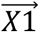 was the unraveled PLS correlation vector (1 x 240; cf. Fig. 2A) and 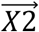 was the unraveled DGE correlation vector (1 x 54; cf. Fig. 5), *s*1^2^ and *s*2^2^were the variances of these vectors, *N*1 was the total number of cross-disease gene module pairs from the latent factor analysis and *N*2 was the number of cross-disease cell type pairs considered in the DGE analysis.

### Identifying biological signaling pathways from gene modules: Gene Ontology enrichment analysis

To query gene co-expression patterns regarding possible underlying biologically meaningful gene programs, we performed a gene set enrichment analysis (GSEA) ^22^. This method is widely used by researchers to gain mechanistic insights into biology based on gene lists derived from omics experiments ^165^. Here, we used the GSEApy python package ^166^, which itself uses Enrichr ^167^. GSEApy is designed to extract statistically over-represented gene sets (example pathways) from a ranked gene list encompassing the whole genome. We focused on the gene ontology (GO) biological process (BP), molecular functions (MF), and cellular component (CC) ^168,169^ databases. Concretely, for each gene module, we fed the gene loadings across the entire protein-coding transcriptome recorded in our datasets to the enrichment tool (*gseapy.Prerank* tool with parameters rnk = gene loadings, min_size = 15, max_size = 1500, and permutation = 1000 for significance testing). We reported the pathways that had a FDR threshold of at most 0.05. This step was repeated identically and independently for all gene modules in each cell across all datasets.

To further verify that the gene set enrichment results were not an artifact of noise in the PLS modeling but rather had actual biological relevance, we turned to our permutation test-derived gene modules. In 1000 permutation iterations, we destroyed the relation between gene expression and disease label, and the thus extracted gene modules captured noise. We fed the gene loadings from these modules into our GSEA pipeline to verify the specificity of the empirical enriched terms.

### Gene network visualization

GO terms are organized in the form of a hierarchical tree which can be downloaded here (OBO 1.4). This hierarchical organization often results in hundreds of hits from an enrichment analysis. One of the techniques widely used to crunch down this dense information is via network visualization ^165,170^. This technique can help identify broad groupings of terms based on a chosen parameter of interest, for example, shared genes between different terms. We utilized Cytoscape ^171^ to create a structured network of disease-relevant GO biological processes identified by an enrichment analysis (cf. above). Each node in the network represented a GO BP hit. The edges were formed based on predefined relationships between nodes conditioned on shared genes and whether they were part of the same regulatory network. The resulting network layout (*yFiles.organic* layout) automatically clustered the enriched terms into biologically meaningful groups, allowing us to identify major functional themes that were shared between AD and PD.

### Differential gene co-expression network

As a complementary analytical pipeline, we sought to explore the transcription profile of the RNA-seq datasets in a top-down gene co-expression network (GCN) approach. Specifically, we started with known disease risk genes from AD or PD GWAS. Using these genes as seeds, we created networks of correlated genes, that is, we identified groups of genes whose differential expression change between disease and control closely matched a seed gene. Seeded differential gene co-expression networks (DGCN) have been previously used to identify regulatory changes in gene expressions across various conditions ^53,55^. At its core, seeded GCNs capture how the gene expressions across the transcriptome are related to the transcription of the seeding gene. By taking a contrastive approach between disease and control (differential), the effects of housekeeping genes are eliminated, and the residual patterns of covariation can be attributed to the effects of a disease. In this study, we used disease-dictated DGCNs to analytically compare the effects of AD or PD on gene expression changes.

To identify our set of seed genes, we searched the most recent GWAS studies that reported AD or PD-associated genes significant at the whole genome level. In doing so, we identified 108 GWAS hits associated with AD ^43^ and 129 GWAS hits associated with PD ^44^. These GWAS hits are the latest nominations in the largest AD or PD-targeted studies. Out of these 237 genes, 5 were common (CTSB, WNT3, BCKDK, HLA-DQA1, and HLA-DRB1) which gave us 232 unique genes. As not all genes are recorded in a snRNA-seq experiment, due to several reasons including technical dropouts or library preparation biases ^172^, we focused on the genes that are present in both AD and PD datasets. We found a total of 164 genes out of the 232 genes that were read in all considered datasets.

Next, for each individual dataset, first, we split them into disease and control groups based on the diagnosis labels provided. We further subdivided each of these groups into subgroups based on cell types. For each cell type *ct*, the co-expression vector for a single GWAS gene *i* with another gene *j* recorded in the snRNA-seq dataset was calculated as *τ*_*b*_(*e*_*i*_, *e*_j_), where *τ*_*b*_ is the Kendall’s tau-b correlation metric, *e*_*i*_ is the read count vector for *i* ∀ observations (nuclei) and *e*_j_is the read count vector of *j* ∀ observations (nuclei). Evaluating *τ*_*b*_across all recorded genes gave us: 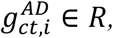 where *M* is the number of genes common to both AD and PD datasets (M = 16,936). Thus, each element of the matrix 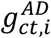 is a numerical value between -1 and 1, capturing the degree of correlation of *j* with GWAS gene *i*. Stacking the vectors for all GWAS genes gave us gene co-expression matrices 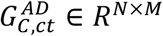 and 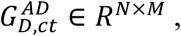 for the control and disease groups respectively, where *N* was the number of GWAS genes (N=164). From this, we formally computed the differential gene co-expression matrix for a single cell type 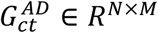 as follows,

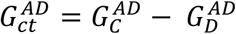

Next, we systematically explored the mutual relationships between the DGCNs across different cell types without discriminating them based on disease. To this end, we employed a hierarchical clustering analysis. Our goal was to probe for clusters of cell types that featured similar genome-wide co-deviation of gene transcription. Concretely, we unraveled 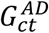 into a vector 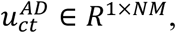 where *NM* = 164 × 16,936 = 2,777,504, and combined them across 6 AD and 9 PD cell types to get *U* ∈ *R*^*C*×*NM*^, where *C* = 15. We computed the linkage matrix based on the Euclidean distance between two unraveled differential co-expression vectors for each cell type (*scipy.cluster.hierarchy.linkage,* parameters method = ‘average’, metric = ‘euclidean’). The linkage algorithm hierarchically clustered the 15 cell types, across AD and PD, with the cluster groups indicating cell types with the closest co-expression patterns. We visualized these clusters as a dendrogram in python (*scipy.cluster.hierarchy.dendrogram*).

We refined our clustering-based qualitative approach to rigorously quantify the association between cross disease DGCNs. Towards this goal, for the *i*th GWAS gene, we computed Kendall’s tau-b correlation metric (*τ*_*b*_) between 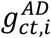 and 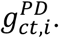 giving us the differential co-expression correlation matrix *G*_*i*_ ∈ *R*^6×9^. Each of these matrices encoded the similarity (-1 to +1; 0 being no association) between AD and PD disease co-expression networks for one gene. To congregate this cross-association information encoded by all the GWAS genes, we vertically stacked the unraveled matrix *G*_*i*_ (unraveled to *g*_*i*_ ∈ *R*^1×54^) into *P* ∈ *R*^164×54^. *P*, in essence, captures multiple modes of information condensed into one matrix: (i) differential gene co-expression between disease and control, (ii) quantified similarity of the expression changes between AD and PD stratified at the level of cell types, (iii) GWAS genes act as seeds not only for the disease in which they were identified but also for other disease. We finally distill this using a classical latent factor method, PCA. This uncovers linear combinations of cross-disease cell types (latent factors) that are most related in terms of their alterations to transcription in response to disease.

The number of significant latent factors that capture biologically meaningful information was determined using a principled permutation testing framework. In 100 permutation iterations, we randomly shuffled the unraveled correlation vector *g*_*i*_, individually for each *i*, thus breaking the inherent meaningful patterns of covariation across cell type pairs. Across the 100 iterations, we fit individual PCA models and computed the explained variances of the derived components. Comparing the permutation variances with our empirical component variances, we retained 4 latent factors as statistically significant based on the 5/95% CI. These four embeddings are by construction uncorrelated and rank-ordered, with the first component capturing the highest amount of variance in *P*.

We conducted a bootstrap analysis on the extracted latent embeddings to formally assess the robustness of the cell type pairs that are closely associated with each other. Across 1000 bootstrap iterations, we sampled different rows (encapsulating all pairwise cell type co-deviations for a GWAS gene) with replacement to simulate a random seed gene collection that could have been sampled from the empirical population. We fit individual PCA models to each of the thus derived samples. To handle the inherent order invariance (changed sequence, especially for later components with small explained variance) and reflection invariance (sign flipping of derived singular vectors) of PCA components, we applied the Jonker-Volgenant algorithm for component matching and Pearson’s correlation (*ρ*) for sign matching. The Jonker-Volgenant algorithm is a widely used technique ^173^ that can identify a one-to-one mapping of latent embeddings derived from two separate bootstrap iterations. This adjustment step was necessary as components that explained similar amounts of variance might swap positions in different sampling runs ^174^. The similarity between a pair of components from two runs was scored using the cosine similarity (cf. above). Subsequently, we solved the optimization problem to maximize the similarity between component orderings from two runs across all pairwise combinations of the first 10 empirical and BS-derived PCA components (*scipy.optimize.linear_sum_assignment*, maximize = True). To align directionality, *ρ* was computed between the empirical PCA component loadings and the BS-component loadings. For cases where *ρ* was less than 1, the latent vector loadings were multiplied by -1. Thus, in a complementary data-driven approach to our latent factor modeling, we could identify combinations of cell types that had the closest associations of gene expression changes between disease and control states.

## Resource Availability

### Data Availability

The snRNA-seq PFC data originated from Mathys, H. et al. Single-cell transcriptomic analysis of Alzheimer’s disease. Nature 570, 332–337 (2019), are available through Synapse (https://www.synapse.org/Synapse:syn18485175) under the doi 10.7303/syn184851755^30^. The data is available under controlled use conditions set by human privacy regulations.

The snRNA-seq MTG data originating from Gabitto, M. I. et al. Integrated multimodal cell atlas of Alzheimer’s disease. Nature Neuroscience, 1–18, is available through SEA-AD consortium’s web portal at SEA-AD.org. Sequencing data are available through controlled access at Sage Bionetworks (accession syn26223298). Sage Bionetworks provide instructions for access to data on the AD Knowledge Portal.

The snRNA-seq substantia nigra data originating from Kamath, T. et al. Single-cell genomic profiling of human dopamine neurons identifies a population that selectively degenerates in Parkinson’s disease. Nature Neuroscience, 25(5) is available from Single Cell Portal (https://singlecell.broadinstitute.org/single_cell/study/SCP1768/)

The snRNA-seq midbrain data originating from Smajić, S. et al. Single-cell sequencing of the human midbrain reveals glial activation and a Parkinson-specific neuronal state. Brain, 145(3) is available for download from the Gene Expression Omnibus (GEO) with accession number GSE157783.

All individual numerical values underlying the summary data presented in the figures, along with the complete code utilized for their generation, will be made openly accessible and deposited in an established open-access repository, such as Zenodo (https://zenodo.org), upon publication of this work.

### Code Availability

Our code will be made available on GitHub at: https://github.com/dblabs-mcgill-mila.

## Acknowledgements

ROSMAP is supported by P30AG10161, P30AG72975, R01AG15819, R01AG17917, U01AG46152, and U01AG61356. ROSMAP resources can be requested at https://www.radc.rush.edu. DB was supported by the Brain Canada Foundation, through the Canada Brain Research Fund, with the financial support of Health Canada, National Institutes of Health (NIH R01 AG068563A, NIH R01 DA053301-01A1, NIH R01 MH129858-01A1), the Canadian Institute of Health Research (CIHR 438531, CIHR 470425), the Healthy Brains Healthy Lives initiative (Canada First Research Excellence fund), the IVADO R3AI initiative (Canada First Research Excellence fund), and by the CIFAR Artificial Intelligence Chairs program (Canada Institute for Advanced Research).

## Author contributions

AB and DB conceptualized the project, planned the experiments and analyzed the results. All authors helped write the manuscript and analyze the results. DB led data analysis.

